# SARS-CoV-2 and MERS-CoV disrupt host protein synthesis via nsp1 with differential effects on the integrated stress response

**DOI:** 10.64898/2025.12.28.696697

**Authors:** Nicholas A. Parenti, Renee Cusic, David M. Renner, Nathaniel Jackson, Chengjin Ye, Li Hui Tan, Jessica J. Pfannenstiel, Anthony R. Fehr, Noam A. Cohen, Luis Martinez-Sobrido, James M. Burke, Susan R. Weiss

## Abstract

Coronaviruses pose a serious threat to public health, driving the need for antiviral therapeutics and vaccines. Therefore, it is paramount to understand how this family of viruses evades cellular antiviral responses and establishes productive infection. The conserved coronavirus non-structural protein (nsp)1 has been shown to inhibit host protein synthesis and promote host mRNA degradation while viral mRNAs are protected. We showed previously that SARS-CoV-2 induces activation of host integrated stress response (ISR) kinases PKR and PERK, which promote phosphorylation of eIF2α and consequent inhibition of host protein synthesis. In contrast, eIF2α remains unphosphorylated during MERS-CoV infection. To investigate the interactions of nsp1 and the ISR kinases, we utilized recombinant SARS-CoV-2 and MERS-CoV expressing nsp1 with mutations in each of two conserved domains. Upon infection with SARS-CoV-2 nsp1 mutants, translation was shut down in wildtype (WT) and PKR knockout (KO) cells but rescued in PERK KO cells, likely due to reduced p-eIF2α. In contrast, translation was rescued during infection with the analogous MERS-CoV nsp1 mutants even in WT cells. Moreover, SARS-CoV-2 WT suppressed expression of GADD34, a negative regulator of eIF2α phosphorylation, while SARS-CoV-2 nsp1 mutants induced GADD34. In contrast MERS-CoV WT induced GADD34. Utilizing single-molecule fluorescence *in situ* hybridization, we found that SARS-CoV-2 and MERS-CoV nsp1 promote host mRNA degradation during WT, but not nsp1 mutant, infection. Finally, while SARS-CoV-2 WT suppressed stress granule formation, nsp1 mutants induced stress granules containing host RNA. Thus, SARS-CoV-2 and MERS-CoV differ in interactions with the ISR and nsp1 control of host protein synthesis.

**Significance:** Coronaviruses cause disease across a wide range of animal species, and the human coronaviruses SARS-CoV-2 and MERS-CoV have caused epidemics of severe respiratory illness. Thus, it is imperative to understand how these viruses antagonize host responses and cause lethal disease. We show here that the betacoronavirus non-structural protein (nsp)1 promotes shutdown of host protein synthesis while preserving viral protein synthesis and, in addition, promotes degradation of host mRNAs. However, SARS-CoV-2 and MERS-CoV differ in their ability to manipulate the host integrated stress response, indicating that it is important to understand detailed coronavirus-host interactions and how they differ even between lethal coronaviruses. Such insights will inform the development of antiviral therapeutics to treat and prevent current and future coronavirus outbreaks.

## Introduction

Pathogenic betacoronaviruses have caused three epidemics in the last approximately twenty years, driving the need for antiviral therapeutics and vaccines. Therefore, it is crucial to understand how this family of viruses evades host cellular antiviral and stress responses and establishes productive infection. We report here our findings on the role of conserved coronavirus non-structural protein 1 (nsp1) during infection with SARS-CoV-2 and MERS-CoV. Nsp1 has been the focus of both *in vitro* and *in cellulo* characterization, but its interaction with host responses during infection remains poorly understood. Moreover, the differences in SARS-CoV-2 and MERS-CoV nsp1-host interaction have not been reported.

Nsp1 has been shown to inhibit translation by binding to the 40S ribosomal subunit at the mRNA entry channel via the conserved C-terminal motif (K164/H165 in SARS-CoV-2 and K181/Y182 in MERS-CoV) (1–4), thereby competitively inhibiting the binding of host mRNAs to the ribosome and preventing translation of host proteins (5). However, viral mRNAs escape nsp1-mediated translation inhibition, due to the leader sequence in the 5’UTR of both genome and subgenomic viral mRNAs (6–9). The viral leader sequence contains RNA secondary structures including several stem loops (SLs) whereby SL1 promotes preferential translation of viral proteins in the presence of nsp1 (1, 3–5, 10, 11).

In addition to its role in inhibiting translation, nsp1 has been shown to mediate the degradation of host, but not viral, mRNAs (12–14) mediated by the conserved N-terminal motif (R124/K125 in SARS-CoV-2 and R146/K147 in MERS-CoV). Similarly, SL1 has been shown to protect viral mRNAs from nsp1-mediated degradation (12, 13, 15). The alanine substitution of the arginine and lysine residues of the nsp1 N-terminal RK motifs of SARS-CoV, MERS-CoV, and SARS-CoV-2 prevents mRNA degradation (4, 10, 15–18). Importantly, it has been shown that the functions of the C– and N-terminal motifs are interdependent whereby mutation of the C-terminal domain disrupts mRNA degradation as well as protein translation initiation, demonstrating that nsp1 must be bound to the ribosome to mediate mRNA cleavage (12, 13, 19). Thus, nsp1 has been shown to disrupt host gene expression via two mechanisms: virus-mediated translation inhibition and mRNA degradation.

The available structural and biochemical data were gathered mainly by overexpression and *in vitro* reconstitution approaches and, thus, do not capture the interactions of nsp1 with host cell responses during viral infection. Thus, it is not clear how the cell responds to the stress of global translation inhibition in the context of infection. Of particular importance is the interaction between nsp1 and the integrated stress response (ISR), two competing mechanisms of translation inhibition (Fig. 1B). We have shown previously that SARS-CoV-2 activates the ISR through at least two kinases: protein kinase R (PKR), which is activated by the accumulation of double-stranded RNA (dsRNA), and PKR-like endoplasmic reticulum (ER) kinase (PERK), which is activated by the accumulation of unfolded proteins in the ER (20, 21). Upon activation, these kinases dimerize, autophosphorylate, and subsequently phosphorylate the alpha subunit of the eukaryotic initiation factor 2 (eIF2α). Phosphorylation of eIF2α results in host-mediated translation inhibition while SARS-CoV-2 protein synthesis continues. In contrast MERS-CoV antagonizes PKR activation (22, 23) and prevents the phosphorylation of eIF2α, despite the activation of the PERK pathway (21). p-eIF2α is detrimental to MERS-CoV protein synthesis (24). Furthermore, GADD34 is transcriptionally upregulated by the ISR and dephosphorylates eIF2α. This rescues protein synthesis and allows the cell to recover from mild ISR stress (25). Importantly, GADD34 expression is also regulated translationally via its upstream open reading frame (uORF), which allows for its preferential expression in the presence of p-eIF2α (26). Because nsp1 disrupts host gene expression via mRNA degradation and translation inhibition, we herein address how GADD34 expression is affected by nsp1, a previously unexplored question.

**Figure 1.**
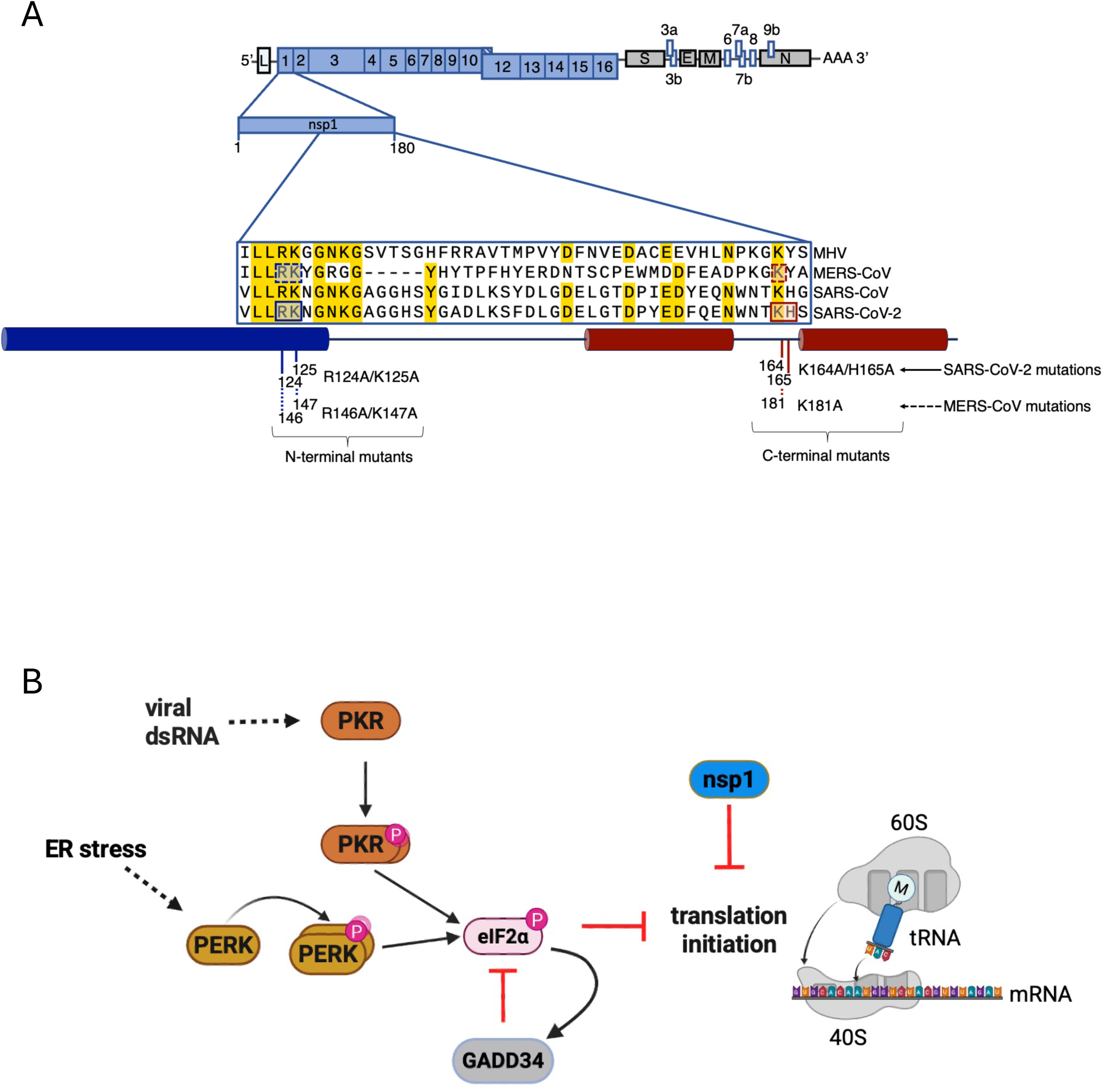
Diagrams of nsp1 motifs in WT and mutant viruses and schematic of integrated stress response. (A) The SARS-CoV-2 genome is shown; the 5’UTR with leader sequence (L), ORFs 1a and 1ab encoded non-structural proteins (nsps) shown in blue and indicated by their number, structural proteins shown in gray: spike (S), envelope (E), membrane (M), and nucleocapsid (N), and accessory proteins shown in white and indicated by their number, and polyA tail (top). Full-length nsp1 of SARS-CoV-2 (middle). Sequence alignment among beta-coronaviruses, from top to bottom: MHV strain A59, MERS-CoV, SARS-CoV, and SARS-CoV-2. N-terminal and C-terminal alanine substitutions are labeled with SARS-CoV-2 amino acid residues boxed in solid lines and sequence numbers under solid lines and MERS-CoV amino acids residues boxed in dashed lines and sequences numbers under dashed lines. (B) Diagram showing the activity of the ISR pathway in response to stress induced by dsRNA or ER stress. Nsp1 is also shown to demonstrate that both p-eIF2α and nsp1 inhibit translation initiation.

The inhibition of global translation by the ISR can lead to the assembly of stress granules, which are large ribonucleoprotein (RNP) complexes that form in the cytosol and contain numerous RNA-binding proteins (G3BP and TIA-1) and translationally stalled mRNAs (27). SGs have been proposed to regulate stress and antiviral responses (28), and SARS-CoV-2 is known to inhibit stress granule formation (14, 29–32). Although the ability of SARS-CoV-2 to inhibit SG formation is attributed to the nucleocapsid (N) protein, by inhibiting the SG-nucleating protein, G3BP1, nsp1 has also been shown to limit the assembly of SGs (33), by a mechanism not yet understood.

To address the role of nsp1 during SARS-CoV-2 and MERS-CoV infection, a reverse genetic approach was utilized to generate recombinant SARS-CoV-2 and MERS-CoV mutant viruses with amino acid substitutions in nsp1 along with CRISPR-Cas9 knock-out (KO) cell lines.

## Results

### Replication of SARS-CoV-2 and MERS-CoV nsp1 mutants is attenuated to different extents in the lung derived A549 cell line and primary nasal epithelial cultures

In order to understand the role of SARS-CoV-2 and MERS-CoV during infection, we generated recombinant viruses with amino acid substitutions in each or both of the key conserved domains of nsp1 (Fig. 1A). For SARS-CoV-2, we generated a C-terminal mutant virus (K164A/H165A or C-term^mut^), an N-terminal mutant virus (R124A/K125A or N-term^mut^), and a double mutant (R124A/K125A/K164A/H165A or C-/N-term^mut^). For MERS-CoV, we used C-terminal (K181A, previously referred to as KA) and N-terminal (R124A/K125A, previously referred to as CD) mutant recombinant viruses, obtained from by Dr. Shinji Makino (16).

We assessed viral replication kinetics in primary nasal epithelial cell air-liquid interface (ALI) cultures (34) or A549 cell lines ectopically overexpressing either angiotensin converting enzyme (ACE2) or dipeptidyl peptidase (DPP4) (20, 22), the receptors for SARS-CoV-2 and MERS-CoV respectively. The three SARS-CoV-2 nsp1 mutant viruses were attenuated in nasal ALI cultures, whereby they exhibited a 10– to 100-fold growth defect compared to wild-type (WT) virus (Fig. 2A). In A549^ACE2^ cells, we observed a smaller yet statistically significant 5-fold growth defect in viral titers of the SARS-CoV-2 nsp1 mutant viruses compared to WT at 24 hours post-infection (24hpi) but recovered to titers comparable to WT by 48hpi (Fig. 2C). Unlike SARS-CoV-2, MERS-CoV nsp1 mutant viruses were not attenuated in nasal ALI cells in comparison to WT virus (Fig. 2B). However, we observed a 10– to 100-fold growth defect of the MERS-CoV nsp1 mutant viruses compared to WT in A549^DPP4^ cells at 24 and 48hpi (Fig. 2D). These data demonstrate that substitution of the residues of nsp1 known to be involved in translation repression and mRNA decay leads to attenuation of viral replication in airway derived cells.

**Figure 2.**
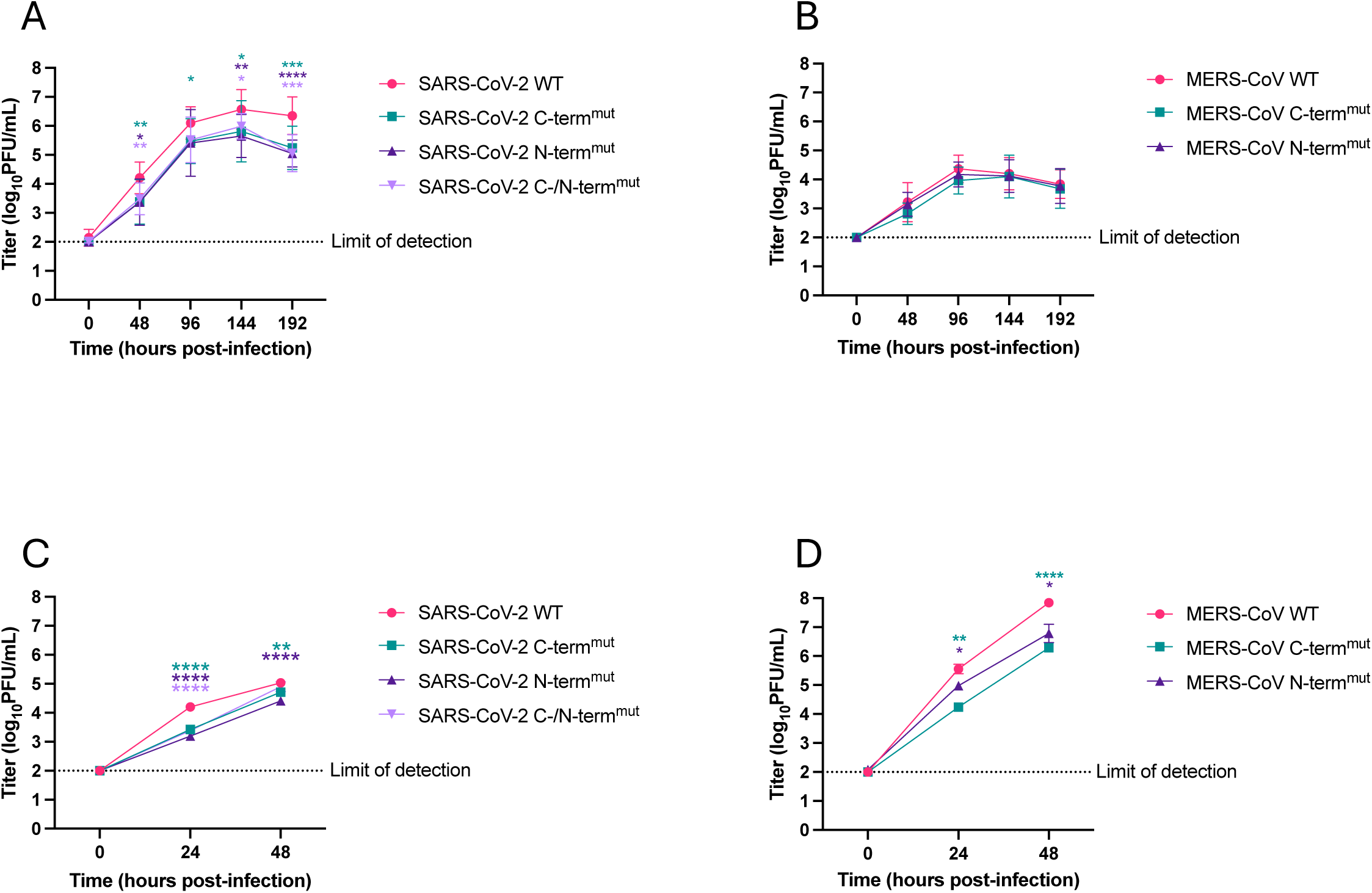
Replication kinetics of SARS-CoV-2 and MERS-CoV recombinant viruses. Cells were infected at MOI=1 PFU/cell (n=3). A) WT SARS-CoV-2 and nsp1 mutant viruses in primary nasal ALI cells. (B) WT MERS-CoV and nsp1 mutant viruses in primary nasal ALI cells. (C) WT SARS-CoV-2 and nsp1 mutant viruses in A549^ACE2^ cells. (D) WT MERS-CoV and nsp1 mutant viruses in A549^DPP4^ cells. Apical fluid (A&B) or supernatants (C&D) were collected at the indicated time post-infection, and viral titers were measured by plaque assay. Data shown is from one of three independent infections (C&D) or average of seven independent experiments (A&B). Values are means ± SD error bars. Statistical significance was determined using a two-way ANOVA. Values are means ± SD error bars. (\**P ≤* 0.05, ** *P ≤* 0.01, *** *P ≤* 0.001, **** *P ≤* 0.0001). Data that are not statistically significant are labeled without significance reported.

### Phosphorylation of eIF2α and inhibition of global translation are differentially regulated during SARS-CoV-2 and MERS-CoV infection

To assess the role of nsp1 in translation inhibition, we performed RiboPuromycylation assays. We observed a reduction in global protein synthesis following infection with WT SARS-CoV-2 in A549 cells (Fig. 3A). However, in contrast to ectopic overexpression studies of nsp1 (1–3), we did not observe a rescue in global protein synthesis during infection with any of the SARS-CoV-2 nsp1 mutant viruses in A549 cells (Fig. 3A). Notably, we observed elevated levels of phosphorylated eIF2α by Western blot analyses, during infection with WT SARS-CoV-2 and all nsp1 mutant viruses (Fig. 3A), which is consistent with previous studies (20, 21, 29). These data suggest that SARS-CoV-2 induces the ISR response, which, together with nsp1, leads to a reduction in bulk protein synthesis. Of note, the electrophoretic mobility of nsp1 during SARS-CoV-2 infection was in agreement with previous studies (1, 2), whereby nsp1 bearing the C-terminal mutation migrates more slowly than WT nsp1.

**Figure 3.**
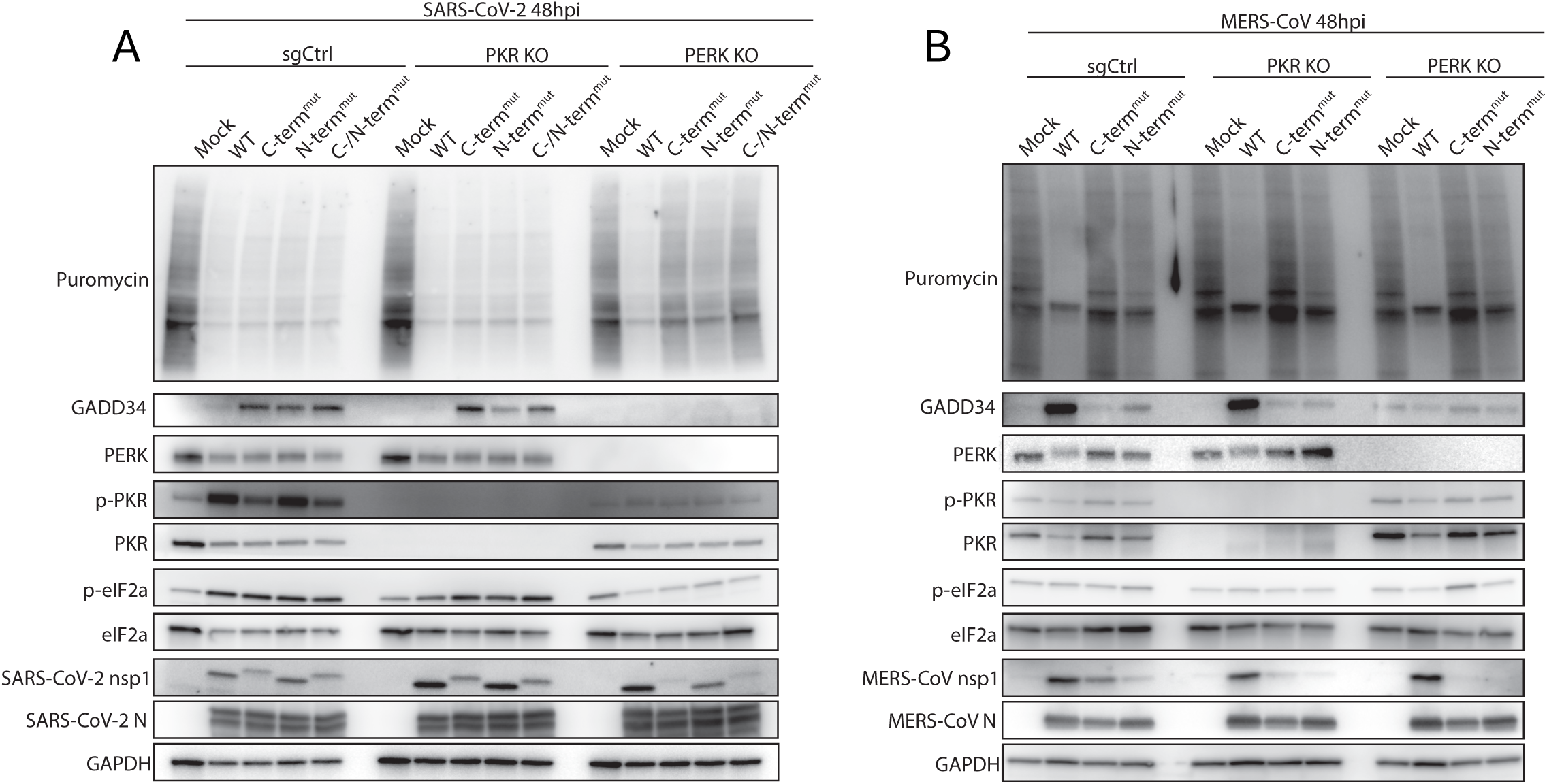
Protein expression in SARS-CoV-2 and MERS-CoV WT and nsp1 mutant infected cells. Cells were infected at MOI=1 PFU/cell. Whole-cell lysates were collected at 48hpi from A549 sgCtrl, PKR KO, or PERK KO cells (expressing either ACE2 for SARS-CoV-2 infections or DPP4 for MERS-CoV infections), mock-infected or infected with (A) SARS-CoV-2 and (B) MERS-CoV WT or nsp1 mutant viruses and analyzed by western blot, probed with antibodies as indicated. In the upper part of panels A&B a RiboPuromycylation assay (55) was used to measure global translation of infected cells, where cells were treated for 10 minutes with 10ug/mL puromycin prior to lysis. Data shown are from one representative of four independent experiments.

Similarly to SARS-CoV-2 infection, we observed a reduction in global protein synthesis following infection with WT MERS-CoV in A549 cells (Fig. 3B). However, global protein synthesis was not reduced following infection with either C-terminal or N-terminal nsp1 mutant viruses. Notably, phosphorylation of eIF2α remained low during infection with all viruses. These data suggest that MERS-CoV nsp1 represses cellular translation by directly inhibiting translation and/or degrading host mRNAs (data shown below) via nsp1 (both C– and N-terminal domains), while eIF2α remains unphosphorylated.

We next examined how the ISR is induced during SARS-CoV-2 infection. We initially considered that it was through activation of PKR, which binds to viral dsRNA and directly phosphorylates eIF2α. However, consistent with our previous findings, eIF2α remains phosphorylated and translation remains shut down during both WT and nsp1 mutant SARS-CoV-2 infection of PKR KO cells (Fig. 3A) (14, 20). We then considered the possibility that SARS-CoV-2 induces the ISR through activation of PERK, which is activated by ER stress and directly phosphorylates eIF2α. Indeed, p-eIF2α levels were reduced in PERK KO cells (Fig. 3A). Furthermore, in the absence of p-eIF2α, we observed a rescue in global protein synthesis during infection with each of the SARS-CoV-2 nsp1 mutant viruses, but not WT SARS-CoV-2, in PERK KO cells (Fig. 3A). Of note, we repeatedly observed an absence of p-PKR in PERK KO cells infected with either WT or nsp1 mutant SARS-CoV-2, which may contribute to the reduction of p-eIF2α expression. Since the lack of PKR activation during infection of the PERK KO cells was surprising, we wanted to be sure these cells were not defective in the ability to activate p-PKR when induced with the synthetic dsRNA poly(I:C). Thus, transfection of poly (I:C) into two independent clones of A549^ACE2^ PERK KO cells, induced robust phosphorylation of both PKR and eIF2α (Fig. S1). Thus, the lack of PKR activation in the PERK KO cells is a consequence of infection that we do not yet understand.

To further understand the differential regulation of the ISR during MERS-CoV and SARS-CoV-2 infection and to probe the roles of viral nsp1 and host p-eIF2α in global translation inhibition, we compared GADD34 protein levels in infected cells. GADD34 is translated from an mRNA that encodes a uORF and, as a result, its expression is upregulated by the ISR (Fig 1B). Thus, we investigated whether GADD34 expression were antagonized by nsp1 in the context of SARS-CoV-2 and MERS-CoV infection. GADD34 protein expression was upregulated during SARS-CoV-2 nsp1 mutant virus infection, but suppressed during WT infection in sgCtrl and PKR KO cells (Fig. 3A), indicating that SARS-CoV-2 antagonizes GADD34 expression in an nsp1-dependent manner. However, GADD34 expression was not observed in SARS-CoV-2 nsp1 mutant-infected PERK KO cells, likely due to the absence of p-eIF2α.

In contrast to SARS-CoV-2, during MERS-CoV infection, eIF2α remains unphosphorylated in sgCtrl, PKR KO, and PERK KO cells. In addition, the global translation phenotypes observed during WT or nsp1 mutant infections were not different in PKR or PERK KO cells compared to WT cells (Fig. 3B). Also, in contrast to SARS-CoV-2, WT MERS-CoV induced strong GADD34 expression (Fig. 3B), consistent with previous findings (21), while MERS-CoV nsp1 mutant viruses do not induce GADD34 expression, likely a consequence of reduced PERK activation in the context of MERS-CoV nsp1 mutant virus infection. Further investigations into the control of GADD34 mRNA and protein expression will be presented and discussed further below.

### SARS-CoV-2 and MERS-CoV nsp1 reduces host mRNAs in the cytosol

SARS-CoV-2 nsp1 has been shown to reduce cytosolic levels of host mRNA (14, 19); however, the precise mechanism by which this occurs remains unsettled. Therefore, we investigated how nsp1 reduces cytosolic host mRNA levels by performing single-molecule fluorescence *in situ* hybridization (smFISH) for human *GAPDH* mRNA following infection with either WT or nsp1 N-terminal and C-terminal mutants of SARS-CoV-2 and MERS-CoV. We also included a SARS-CoV-2 mutant in which residues (D33/E36/E37/E41) proposed to interact with nuclear transcription factor, X-box binding 1 (NFX1) to inhibit nuclear mRNA export are substituted (nsp1^D33K/E36K/E37K/E41L^) (35). Notably, we performed these analyses in A549 cells lacking RNase L (RL KO), as activated RNase L can phenocopy nsp1 by degrading cellular RNA (36).

In A549 RL KO cells infected with WT SARS-CoV-2, we observed a robust loss (10– and 100-fold reduction) in *GAPDH* mRNA molecules at 24hpi in comparison to mock-infected cells (Fig. 4A-B). We observed a loss in *GAPDH* mRNA as early as 12hpi (Fig. 4B; Fig. S2A), and *GAPDH* mRNA levels further reduced by 48hpi (Fig. 4B; Fig. S2B). In. contrast to WT SARS-CoV-2, we did not observe a loss in *GAPDH* mRNA in cells infected with the nsp1 N-terminal and C-terminal mutant viruses (Fig. 4A,B, Fig. S2A,B), nor did we observe the loss of *GAPDH* mRNA in cells infected with the SARS-CoV-2 nsp1^D33K/E36K/E37K/E41L^ mutant (Fig. 4A,B). Notably, we observed similar results in WT A549 cells, though a large fraction of cells infected with the nsp1 mutant viruses display a loss of *GAPDH* mRNA by 48hpi (Fig. S3A-D). Because RNase L activates between 24 and 48hpi in response to SARS-CoV-2 (14, 20, 23), and because the loss of *GAPDH* mRNA did not occur in RL KO cells infected with nsp1 mutants, these data suggest that the loss of cellular mRNA late during infection in WT A549 cells infected with the nsp1 mutants is the result of RNase L-mediated mRNA decay. Combined, these data show that residues involved in translation inhibition (C-terminal, N-terminal), as well as the NFX1-interacting residues (D33,E36,E37,E41), are required nsp1-mediated reduction of cellular mRNA in the cytosol.

**Figure 4.**
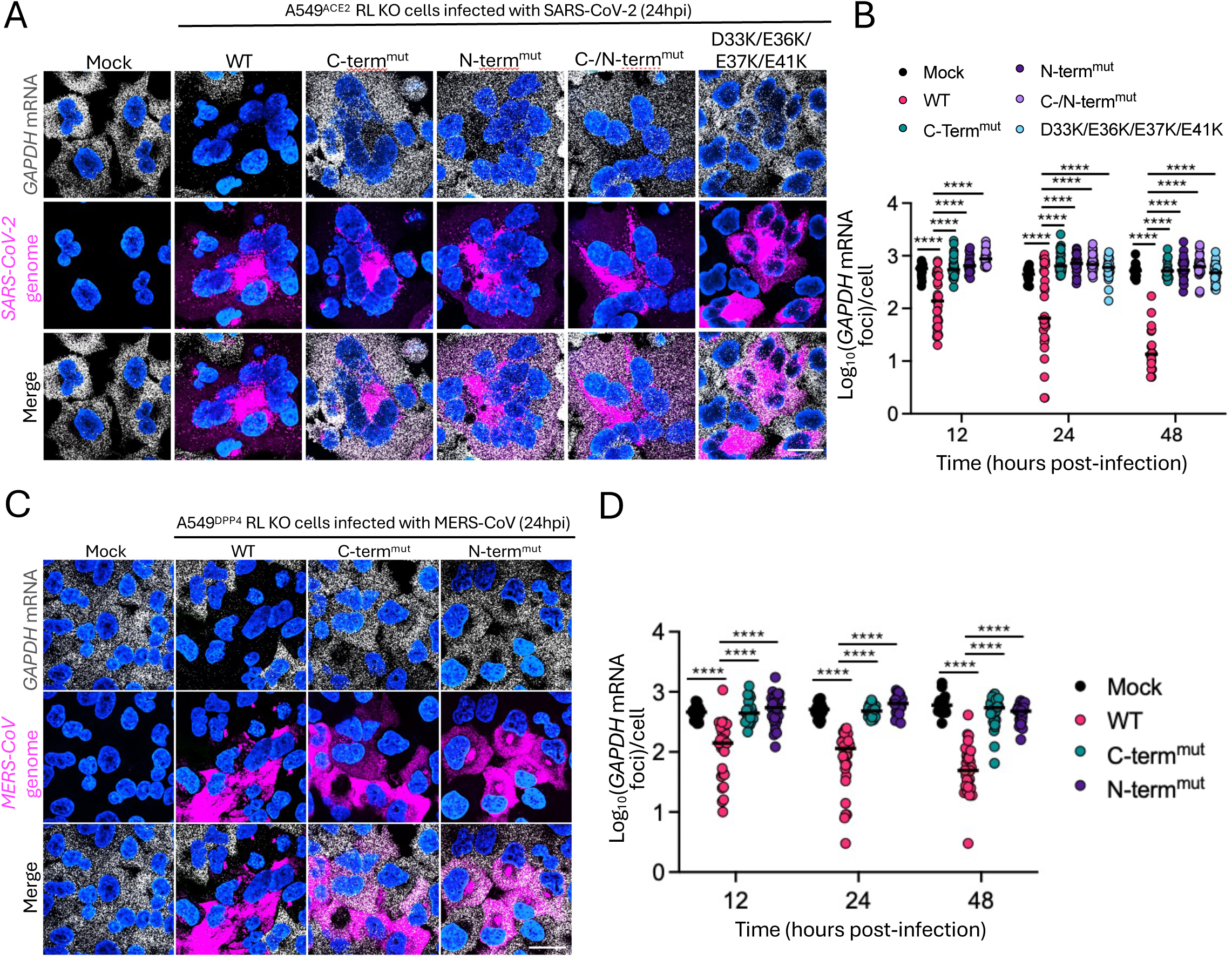
SARS-CoV-2 and MERS-CoV nsp1 is required for viral-mediated degradation of cellular mRNA. (A) smFISH for SARS-CoV-2 genome RNA (ORF1a) and human *GAPDH* mRNA in A549^ACE2^ RL KO cells infected with WT or nsp1 mutant SARS-CoV-2 viruses at 24hpi. Images are representative of 3 replicate experiments. (B) Quantification of *GAPDH* mRNA in individual cells (dots) at indicated time post-infection as represented in (A). Minimum of 16 cells per viral infection were quantified from 3 or more fields of view. Quantification is representative of 3 biological replicates.(C) smFISH for MERS-CoV genome RNA (ORF1a) and human *GAPDH* mRNA in A549^DPP4^ RL KO cells infected with WT or nsp1 mutant MERS-CoV viruses at 24hpi. Images are representative of 3 biological replicates. (D) Quantification of *GAPDH* mRNA in individual cells (dots) at indicated time post-infection as represented in (C). Minimum of 22 cells were quantified per virus from 2 or more fields of view. Quantification is representative of 3 biological replicates.Scale bars represent 20μm.

Similarly, cells infected with WT MERS-CoV displayed a significant (>5-fold) reduction in *GAPDH* mRNA molecules by 24hpi comparison to mock-infected cells (Fig. 4C,D). Remarkably, we observed this robust reduction in *GAPDH* mRNA as early as 12hpi (Fig. 4D; Fig. S5A), and a further reduction (10– to 100-fold) by 48hpi (Fig. 4D; Fig S5B). In contrast, cells infected with either N-terminal or C-terminal nsp1 mutant viruses contained significantly higher *GAPDH* mRNA levels over the course of infection, whereby *GAPDH* mRNA levels remained comparable to the levels observed in mock-infected cells (Fig. 4C,D; Fig. S4A,B). Unlike SARS-CoV-2, MERS-CoV does not activate RNase L (22, 23), we observed similar results in WT and RNase L KO A549 cells (Fig. S4C-F). Because the N-terminal and C-terminal residues are required for mRNA degradation, these data are in agreement with previous *in vitro* studies, which suggest that nsp1 must be bound to the ribosome via its C-terminal domain in order to mediate mRNA degradation (12, 13).

### Nsp1 reduces host mRNAs in the cytosol via accelerated decay

We next investigated how nsp1 reduces cytosolic cellular mRNA levels. Two observations support that SARS-CoV-2 nsp1 accelerates the decay of mRNA in the cytosol. First, although SARS-CoV-2 nsp1 has been proposed to inhibit mRNA export by interacting with NFX1 (35), we did not observe an increase in the absolute number of *GAPDH* mRNA molecules in the nucleus of cells infected with either WT SARS-CoV-2 or WT MERS-CoV in comparison to mock-infected cells (Fig. 5A-D). Moreover, we did not observe less *GAPDH* mRNA in the nucleus of cells infected with the SARS-CoV-2 nsp1^D33K/E36K/E37K/E41L^ mutant in comparison to WT SARS-CoV-2. Infections with the C-terminal and N-terminal nsp1 mutants of SARS-CoV-2 and MERS-CoV, which do not lead to a loss of cytosolic *GAPDH* mRNA, also did not promote reduced nuclear *GAPDH* mRNA levels compared the WT viruses.

**Figure 5.**
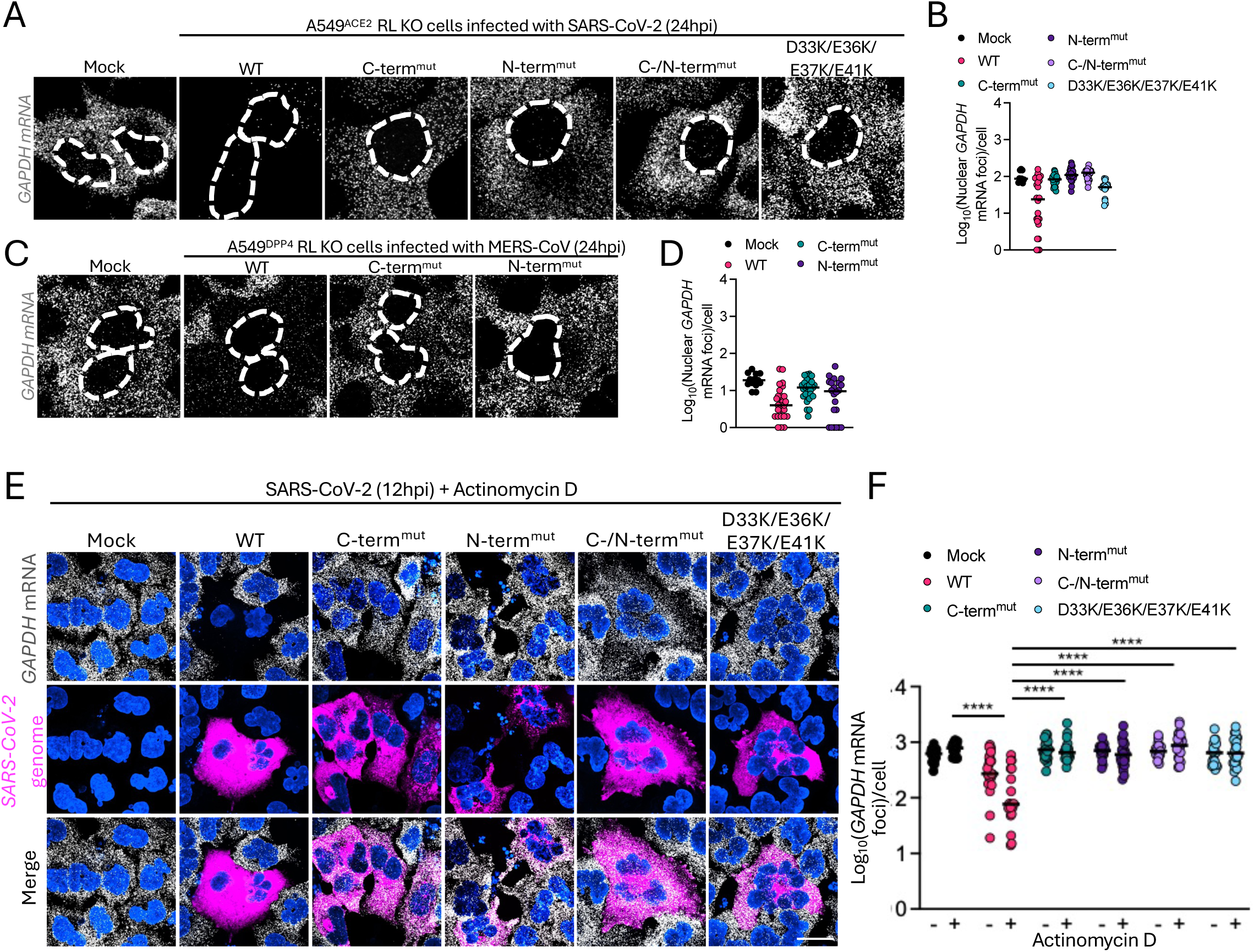
SARS-CoV-2 and MER-CoV nsp1 do not cause mRNA nuclear export block. (A) smFISH for human *GAPDH* mRNA in A549^ACE2^ RL KO cells infected with WT or nsp1 mutant SARS-CoV-2 viruses at 24hpi. Images are representative of 3 biological replicates.(B) Quantification of nuclear *GAPDH* mRNA in individual cells (dots) for indicated viruses as represented in (A). Minimum of 10 cells quantified per virus, quantification is representative of 3 biological replicates. (C) smFISH for human *GAPDH* mRNA in A549^DPP4^ RL KO cells infected with WT or nsp1 mutant MERS-CoV viruses at 24hpi. Images are representative of 3 biological replicates. (D) Quantification of nuclear *GAPDH* mRNA in individual cells (dots) for indicated viruses as represented in (C). Minimum of 10 cells quantified per virus, quantification is representative of 3 biological replicates. (D) smFISH for SARS-CoV-2 genome RNA (ORF1a) and human *GAPDH* mRNA in A549^ACE2^ RL KO cells infected with WT or nsp1 mutant SARS-CoV-2 viruses at 12hpi with 1mg/mL of Actinomycin D. Images are representative of 3 or more fields of view. (E) Quantification of *GAPDH* mRNA in individual cells (dots) for indicated viruses with or without 1mg/mL Actinomycin D treatment as represented in (D). Minimum of 18 cells from a minimum of 3 fields of view quantified per virus. Scale bars represent 20μm.

Second, we tested if treatment with actinomycin D, which inhibits transcription and would thus phenocopy the cytosolic loss in *GAPDH* mRNA via a potential nsp1-mediated nuclear export block, would lead to a reduction in GAPDH mRNA at 12hpi. Importantly, we did not observe a reduction in *GAPDH* mRNA in cells infected with the SARS-CoV-2 nsp1 mutant viruses treated with actinomycin D, whereas GAPDH mRNA was reduced in cells infected with WT SARS-CoV-2 (Fig. 5E,F). We obtained a similar result with MERS-CoV, whereby GAPDH mRNA remained abundant despite treatment with actinomycin D (Fig. S5A,B). We also confirmed that actinomycin D inhibited transcription by demonstrating that interferon-beta induction in response to poly(I:C) was abolished in cells treated with actinomycin D (Fig. S5C). These data show that neither transcription nor nuclear mRNA export are required to maintain cytosolic *GAPDH* mRNA levels over the course of infection in cells infected with nsp1 mutant viruses. Thus, *GAPDH* mRNA is stable in the cytosol of cells infected with SARS-CoV-2 nsp1 mutant viruses, whereas *GAPDH* mRNA is unstable in cells infected with SARS-CoV-2 encoding a WT nsp1.

Combined, these data show that SARS-CoV-2 and MERS-CoV nsp1 promote reduced levels of cellular mRNA in the cytosol by accelerating mRNA decay, and that the N-terminal and C-terminal residues are required for this effect. Lastly, these data show that D33/E36/E37/E41 residues are also required for nsp1 to accelerate the decay rate of cellular mRNAs in the cytosol.

### GADD34 evades nsp1-mediated mRNA decay, but not translation repression in SARS-CoV-2 infected cells

We found above that SARS-CoV-2 induces the ISR via PERK-mediated phosphorylation of eIF2α. Furthermore, we found that GADD34, an ISR induced protein, was expressed in cells infected with SARS-CoV-2 nsp1 mutant viruses, but not WT (Fig. 3A). Since GADD34 regulates translation through p-eIF2α dephosphorylation, we sought to further understand the expression of GADD34 and its absence in cells infected by WT SARS-CoV-2 (Fig. 3). Immunofluorescence assays (IFAs) confirmed that GADD34 protein synthesis was rarely induced in cells infected with WT SARS-CoV-2, whereas cells infected with the C– or N-terminal nsp1 mutant viruses frequently induced GADD34 protein in comparison to mock-infected cells (Fig. 6A,B). These data indicate that nsp1 inhibits the induction of GADD34 protein.

**Figure 6.**
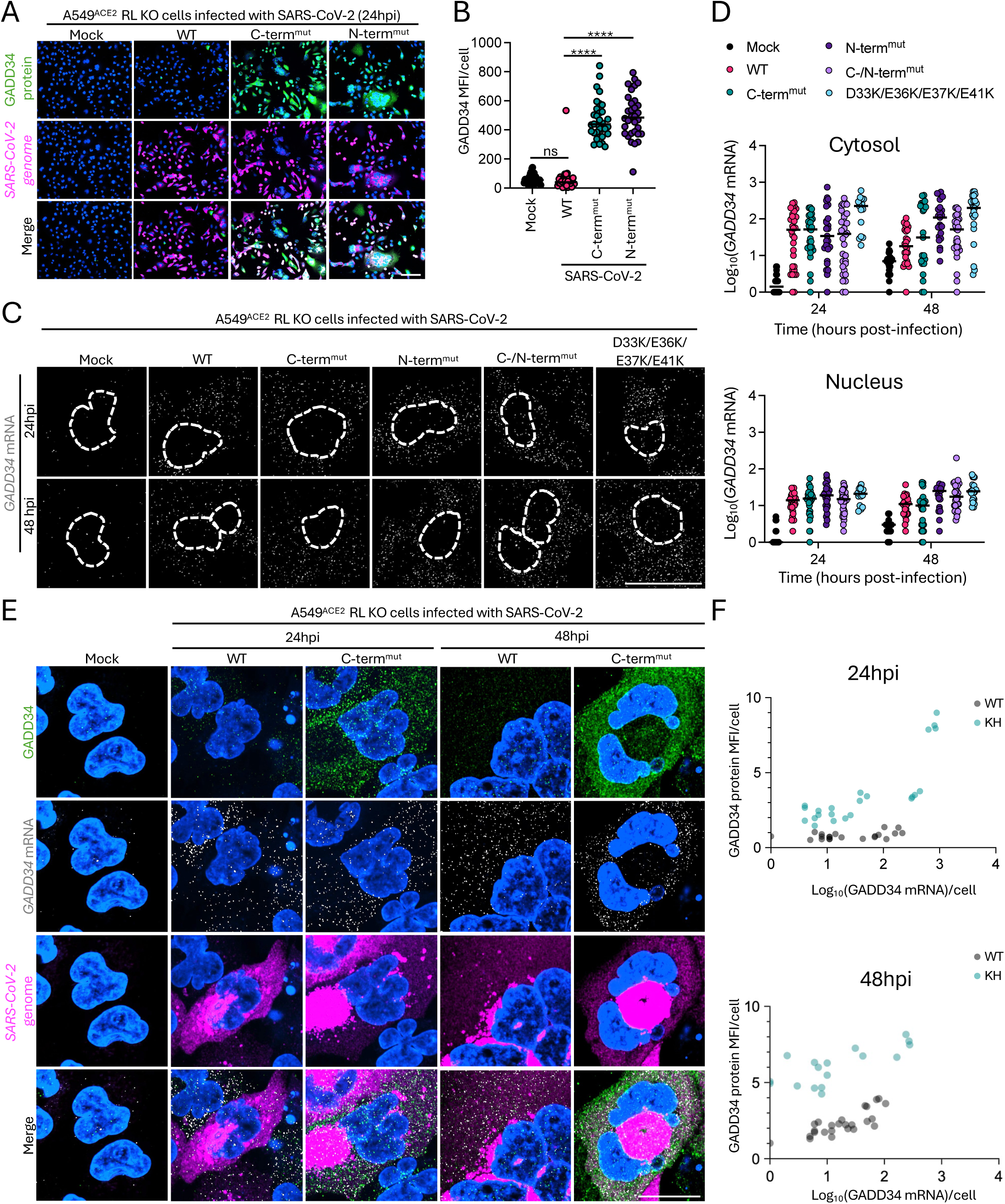
SARS-CoV-2 nsp1 suppresses GADD34 expression via translation inhibition. (A) IFA analyses for GADD34 protein and smFISH for SARS-CoV-2 genome RNA (ORF1a) in A549^ACE2^ RL KO cells infected with WT or nsp1 mutant SARS-CoV-2 viruses at 24hpi. Images representative of 4 biological replicates. (B) Quantification of GADD34 mean fluorescence intensity (MFI) in individual cells (dots) for viruses as represented in (A). Minimum of 30 cells quantified from 2 or more fields of view per virus. Quantification is represenetative of 4 biological replicates. (C) smFISH for human *GADD34* mRNA in A549^ACE2^ RL KO cells infected with WT or nsp1 mutant SARS-CoV-2 viruses at indicated times post-infection. Images representative of 4 biological replicates. (D) Quantification of nuclear and cytosolic *GADD34* mRNA in individual cells (dots) at indicated time post-infection for indicated viruses as represented in (C). Minimum of 15 cells were quantified per virus, quantification is representative of 4 biological replicates. (E) IFA analyses for GADD34 protein and smFISH for SARS-CoV-2 genome RNA (ORF1a) and human *GADD34* mRNA in A549^ACE2^ RL KO cells infected with WT or nsp1 mutant SARS-CoV-2 viruses at indicated times post-infection. Images representative of 4 biological replicates. (F) Quantification of GADD34 MFI and of *GADD34* mRNA in individual cells (dots) for indicated viruses at indicated times post-infection as represented in (E). Minimum of 20 cells quantified per virus, quantification representative of 4 replicates. Scale bars represent 20μm.

To determine the precise mechanism by which SARS-CoV-2 nsp1 inhibits GADD34 protein expression, we examined *GADD34* mRNA induction and localization by smFISH at 24 and 48hpi. We observed that *GADD34* mRNA was induced in response to WT SARS-CoV-2 infection as well as infection with each of the nsp1 mutants in comparison to mock-infected cells (Fig. 6C,D). We observed that *GADD34* mRNA was primarily localized to the cytosol in cells infected with WT or nsp1 mutants, whereby ∼100 *GADD34* mRNAs were localized in the cytosol and ∼10 mRNAs were localized in the nucleus (Fig. 6C,D). Importantly, *GADD34* mRNA levels in both the cytosol and nucleus were comparable in cells infected with WT SARS-CoV-2 and the nsp1 mutants at 24hpi (Fig. 6C,D), though cytosolic *GADD34* mRNA reduced in cells infected with WT virus to levels lower than those observed in cells infected with nsp1 mutants at 48hpi. Combined, these data indicate that nsp1 does not inhibit the export of GADD34 mRNA from the nucleus, nor robustly accelerate its decay. Instead, these data suggest that nsp1 inhibits the translation of the GADD34 mRNA.

To determine if nsp1 indeed represses the translation of *GADD34* mRNA, we performed IFA and smFISH analyses for GADD34 protein and *GADD34* mRNA, respectively, in cells infected with either WT SARS-CoV-2 or C-terminal nsp1 mutant virus. These analyses showed that GADD34 protein levels were significantly lower in cells infected with WT virus in comparison to the C-terminal nsp1 mutant virus despite similar *GADD34* mRNA induction (Fig. 6E,F). Combined, these data suggest that *GADD34* mRNA is translationally repressed by nsp1, but its decay rate is not drastically increased by nsp1, suggesting that *GADD34* mRNA evades nsp1-mediated mRNA decay. Moreover, our data show that the RK, KH, and N-terminal D33/E36/E37/E41 residues are required to inhibit the translation of the *GADD34* mRNA. Unlike SARS-coV-2, we observed that GADD34 was induced in cells infected by WT but not nsp1 mutants. We do not understand why MERS-CoV nsp1 differs from SARS-CoV-2 in regards to GADD34 induction, but we are currently investigating this interesting and ciritical difference.

### Nsp1 inhibits stress granule assembly

Our above data show that SARS-CoV-2 nsp1 suppresses cellular translation, inhibits PERK-mediated ISR, limits GADD34 protein synthesis, and accelerates decay of cellular mRNA. Because these functions could alter the assembly of SGs, we examined how SARS-CoV-2 nsp1 alters SGs by performing IFA for three SG markers, poly(A)+RNA, TIA-1, and G3BP1. In mock infected cells, poly(A)+RNA, TIA-1 and G3BP1 were diffusely localized in the cytosol. In cells infected with WT SARS-CoV-2, poly(A)+RNA and TIA1 assembled into small punctate granules (Fig. 7A,B).

**Figure 7.**
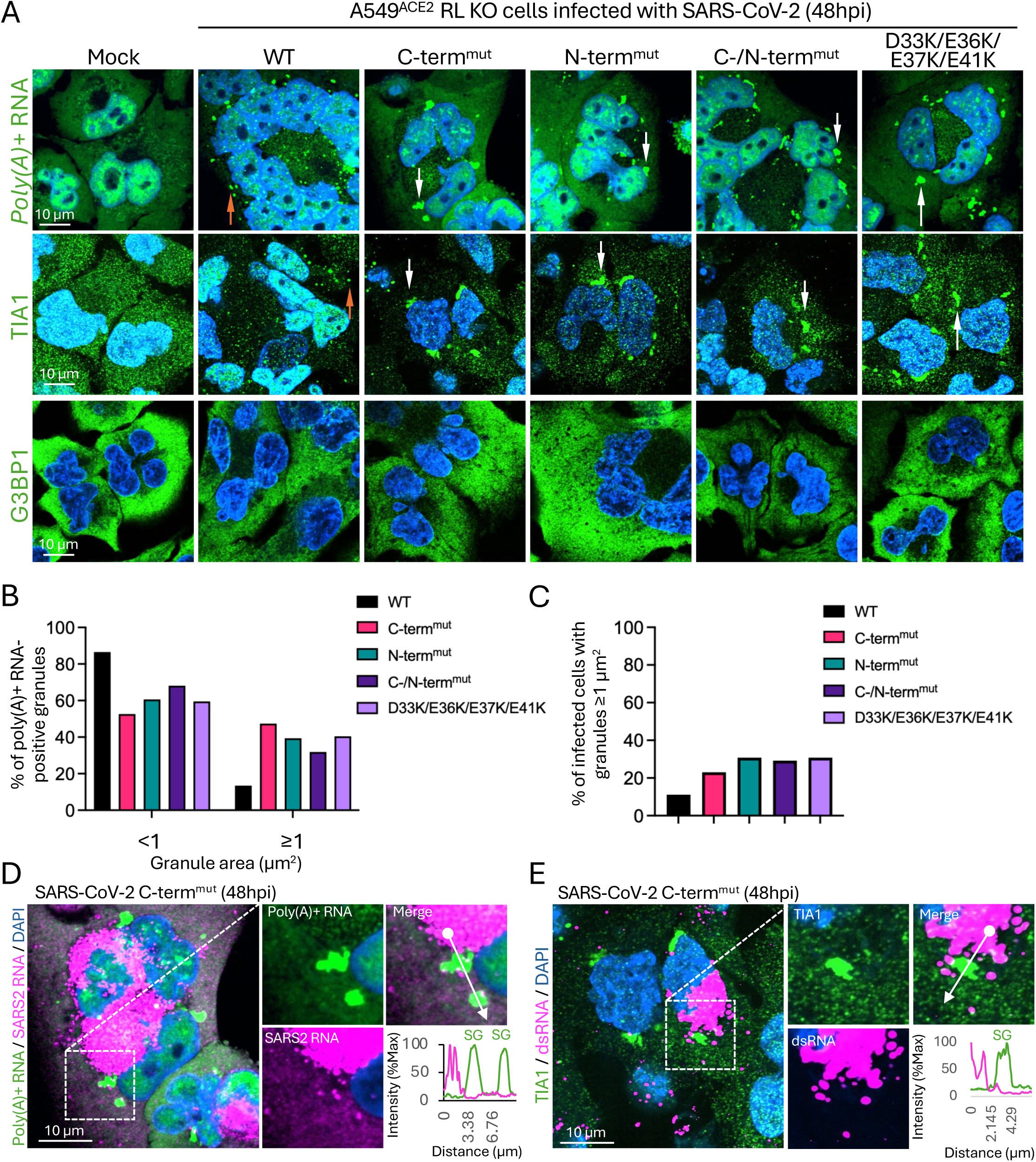
SARS-CoV-2 nsp1 limits stress granule assembly. (A) IFA analyses for stress granule markers, TIA-1 and G3BP1, and FISH for poly(A)+RNA 48 hours post-infection with WT or nsp1 mutant SARS-CoV-2 viruses in A549^ACE2^ RNase L-KO (RL-KO) cells. Images representative of 4 or more fields of view. (B) The percentage of FISH poly(A)+RNA granules smaller or larger than 1 μm^2^ in cells infected with WT or nsp1 mutants. Minimum of 24 cells were quantified per virus. (C) Histogram displaying the percentage of cells with poly(A)+RNA granules equal to or larger than 1 μm^2^. Minimum of 24 cells quantified per virus. (D) smRNA-FISH for SARS-CoV-2 genome RNA (ORF1a) and poly(A)+RNA. The graph displays the normalized fluorescence intensity (% of maximum intensity) of poly(A)+RNA and SARS-CoV-2 genome RNA along the line in the merged inset image panel. Images representative of 4 or more fields of view. (E) Similar to (D) but IFA for TIA1 and dsRNA. Scale bars represent 10μm.

Most strikingly, we observed that cells infected with the SARS-CoV-2 nsp1 mutants contained large granules composed of poly(A)+RNA and TIA-1 (Fig. 7A). The assembly of these larger granules was more in frequent in cells infected with nsp1 mutants compared to WT SARS-CoV-2 (Fig. 7B and C). The large size and granular morphology of these granules is consistent with them being SGs. However, we note that the SGs lack G3BP1, which is presumably due to N-mediated inhibition of G3BP1 (further addressed in the discussion).

We next asked if the SGs that form in cells infected with the SARS-CoV-2 nsp1 mutants contain viral RNA or dsRNA. To determine if the SGs contained SARS-CoV-2 genome RNA, we performed smRNA-FISH for SARS-CoV-2 genome sequences. We co-stained cells for poly(A)+RNA. Importantly, we did not observe enrichment of SARS-CoV-2 genome RNA in the poly(A)+RNA-positive SGs (Fig. 7D). Similarly, co-IFA for dsRNA and TIA-1 showed that the TIA1-positive SGs did not enrich for dsRNA (Fig. 7E). These data indicate that the SGs that assemble in cells infected with the SARS-CoV-2 nsp1 mutants are composed of cellular mRNA. Combined, these data indicate that SARS-CoV-2 nsp1 limits the assembly of stress granules.

## DISCUSSION

Coronavirus nsp1 has been the subject of many studies conducted mostly via ectopic overexpression. These studies have shown that nsp1 of SARS-CoV (4, 11, 37, 38), MERS-CoV (3, 15, 16), and SARS-CoV-2 (1, 2, 5, 12, 13, 19, 39–42) bind to ribosomes and inhibit translation and in addition promote degradation of host mRNAs. Some studies of recombinant nsp1 mutant viruses of SARS-CoV (18, 38), MERS-CoV (15), and SARS-CoV-2 (17) recapitulate the in vitro findings of protein synthesis inhibition and mRNA decay and demonstrate that WT nsp1 is necessary for optimal replication. However, none of these studies address how nsp1 interacts with the host suppression of protein synthesis in response to virus infection through the ISR, the focus of our study reported here.

We investigated the function of nsp1 during infection with two lethal coronaviruses, SARS-CoV-2 and MERS-CoV, with a focus on how nsp1 interacts with the host ISR pathway in the context of bona fide infection. We compared recombinant mutant viruses of SARS-CoV-2 and MERS-CoV containing amino acid substitutions in conserved nsp1 domains to their WT counterparts. Both SARS-CoV-2 and MERS-CoV nsp1 mutant viruses are replication deficient compared to their respective WT counterparts, but this varies between infections of the A549 cell line and primary nasal epithelial ALI cultures (Fig. 2). Furthermore, we found that both SARS-CoV-2 and MERS-CoV inhibit host cell translation and mediate decay of cellular mRNA via nsp1, similar to findings in studies in which nsp1 is overexpressed (1–3, 5, 12, 13, 15, 16, 19, 39–42). In contrast, SARS-CoV-2 and MERS-CoV, behave quite differently in their interactions with the ISR, notably in the expression of p-eIF2α and GADD34 as discussed below.

We found that, during SARS-CoV-2 infection, host translation is inhibited both by nsp1 and p-eIF2α. Infection with SARS-CoV-2 WT activates at least two ISR kinases, PKR and PERK, which lead to phosphorylation of eIF2α and host-mediated translation inhibition. Consequently, mutation of nsp1 alone does not rescue global translation. We show that global translation is rescued during nsp1 mutant virus infection only in the absence of PERK expression. Importantly, PERK KO cells infected with nsp1 mutant viruses fail to phosphorylate eIF2α above mock-infected cells. Conversely, PKR KO cells infected with either WT or nsp1 mutant viruses phosphorylate eIF2α at levels equivalent to infected control (sgCtrl) cells. This demonstrates that PERK activation drives greater eIF2α phosphorylation compared to PKR during SARS-CoV-2 infection. This indicates that PERK is the major contributor to eIF2α phosphorylation during SARS-CoV-2 infection and, thus, host-mediated translation inhibition. Furthermore, surprisingly PKR phosphorylation depended on PERK during SARS-CoV-2 infection (Fig 3) but not when induced by the synthetic dsRNA, poly(I:C) (Fig. S1). We speculate that there may be some cooperativity between PERK and PKR during infection, but understanding this finding will require further investigation.

We found that GADD34 protein expression is suppressed by WT SARS-CoV-2 nsp1 (Fig. 3 and Fig. 6). This is noteworthy because GADD34 is translated from an mRNA bearing a uORF, and thus is expected to be translated in the presence of p-eIF2α when the ISR is activated. In contrast, nsp1 mutant viruses induce robust GADD34 protein expression in both control (sgCtrl) and PKR KO cells. Of note, in PERK KO cells, we do not observe GADD34 protein, presumably due to the reduction in eIF2α phosphorylation. Given that GADD34 translation is increased in the presence of p-eIF2α, these data indicates that WT nsp1 shuts down host protein synthesis even of mRNAs bearing a uORF, suggesting that nsp1 directly disrupts the ISR.

In contrast to SARS-CoV-2, MERS-CoV infection does not induce eIF2α phosphorylation. Thus, MERS-CoV relies on nsp1 alone to shutdown protein synthesis. Indeed, MERS-CoV replication appears to be optimized in the absence of p-eIF2α. Inducing the phosphorylation of eIF2α in MERS-CoV infected cells, by either employing an immunostimulatory mutant MERS-nsp15^mut^/ΔNS4a (22, 23, 43) or treating cells with the phosphatase inhibitor salubrinal (24), reduces replication. Nevertheless, we infer that WT MERS-CoV may induce some level of p-eIF2α via PERK, but that GADD34 expression is great enough to reverse this, thus providing for the optimal translational environment for MERS-CoV. Importantly, there is no detectable GADD34 expression in the context of WT MERS-CoV infection in PERK KO cells, supporting the argument that PERK-mediated ISR activation drives GADD34 expression during MERS-CoV infection. Furthermore, we show that adequate PERK activation and subsequent GADD34 expression depends on nsp1 as mutant viruses do not induce GADD34. Thus, we provide evidence that, in the context of MERS-CoV infection, nsp1 is a major contributor to ER stress, subsequent PERK activation, and ultimately GADD34 expression.

As discussed above our data highlight striking differences between SARS-CoV-2 and MERS-CoV infections as to their interactions with the ISR. Our data support the notion that SARS-CoV-2 nsp1 is more efficient in shutting down protein synthesis than MERS-CoV nsp1. Indeed, biochemical data show that SARS-CoV-2 nsp1 binds to the mRNA entry site of the 40S ribosome with greater affinity than MERS-CoV nsp1 (3). Moreover, MERS-CoV nsp1 had a lower binding affinity than that of bat-Hp-CoV (3), which like SARS-CoV-2 nsp1, has a KH amino acid sequence in the C terminal domain, while MERS nsp1 has a KY sequence (Fig. 1). It is possible that the greater binding affinity of SARS-CoV-2 nsp1 allows for more complete suppression of host mRNA translation including suppression of GADD34 and potentially other ISR related proteins bearing a uORF. This may allow for SARS-CoV-2 to scavenge the available non-phosphorylated eIF2α within the cell more effectively and allow translation of viral proteins in the presence of p-eIF2α. Indeed SARS-CoV-2 WT replicates to high titer even in the presence of p-eIF2α (Fig. 2 and Fig. 3).

Nsp1 was shown by several groups to promote reduced levels of cytoplasmic host mRNA (19, 35). This has been proposed to occur by inhibiting nuclear mRNA export (35), and/or by accelerating degradation of host mRNA either by functioning as a nuclease or inducing a host nucleolytic activity (11–13, 16, 44). While the precise mechanism is still poorly understood purified nsp1 combined with host mRNA does not cause mRNA degradation consistent with the absence of a motif indicative of a nuclease (12, 13). However, nsp1 combined with host mRNA and purified 40S ribosomes does promote mRNA degradation (12, 13, 19). Consistent with these reports, our smFISH data show that while WT viruses do mediate a robust reduction in *GAPDH* mRNA, both the N-terminal and C-terminal mutants as well as the D33,E36,E37,E41 nsp1 mutants fail to promote degradation of host mRNA Fig 4 and Fig. S2-4). The quantification of host *GAPDH* mRNA following infection with WT or nsp1 mutants, combined with actinomycin D treatment (Fig. 5 and Fig. S5), demonstrates that nsp1 reduces cellular mRNA levels by accelerating the decay of bluk cellular mRNA in the cytosol. Although it is possible that cytoplasmic mRNA degradation can lead to an mRNA export block as reported previously (14) our observations do not provide evidence supporting that nsp1 inhibits the export of *GAPDH* mRNA or *GADD34* mRNA during WT SARS-CoV-2 infection, thus warranting further investigation into the role of nsp1 in regulating host mRNA export.

We assessed the effects of SARS-CoV-2 nsp1 on host mRNA translation apart from its activity of mRNA degradation. Expression of most host proteins, for example GAPDH, will be suppressed by nsp1 mediated mRNA degradation, which would confound our measurement of nsp1-mediated translation suppression. Thus, we monitored GADD34 expression, which is translated from an mRNA with a uORF in the presence of p-eIF2α (26). Our data show that GADD34 protein is significantly repressed by nsp1 by western blot (Fig. A) and IFA (Fig. 6A,B). However, GADD34 mRNAs were induced to equivalent levels in cells infected with WT or nsp1-mutant SARS-CoV-2 at 24hpi, and were only slightly reduced by nsp1 by 48-hpi (Fig. 6C). This suggests that *GADD34* mRNA evades nsp1-mediated decay, and thus nsp1 represses translation of *GADD34* mRNA. Indeed, staining for both *GADD34* mRNA and GADD34 protein in WT and nsp1 mutant virus infection demonstrated that GADD34 protein but not mRNA is reduced during SARS-CoV-2 WT infection (Fig. 6). Therefore, we conclude that SARS-CoV-2 nsp1 suppresses GADD34 protein levels via translation inhibition of the *GADD34* mRNA. Taken together with the evidence that nsp1 suppresses GAPDH via accelerated mRNA degradation, these data provide evidence that SARS-CoV-2 nsp1 redundantly antagonizes host gene expression. We do not understand why *GAPDH* mRNA is degraded while *GADD34* mRNA is not; this feature of nsp1-mediate mRNA degradation warrants further investigation. However, one study shows that *GADD34* mRNA contains an AU-rich element in its 3’UTR, which is recognized by the ZFP36 family of proteins and promotes its stabilization during cellular stress (45), and cellular transcripts containing 5′ terminal oligopyrimidine tracts (5′-TOP RNAs) and the TIAR mRNA have been shown to evade nsp1-mediated translation and/or mRNA decay (46)(33). Future work will investigate how the GADD34 mRNA transcript evades nsp1-mediated decay.

Our data show that SARS-CoV-2 nsp1 limits the formation of stress granules. This is based on the observation that all the nsp1 mutant viruses induced the assembly of large cytoplasmic granules that contained poly(A)+RNA and TIA-1 (Fig. 7A-C). In contrast, cells infected with WT SARS-CoV-2 assembled smaller punctate granules that resemble RNase L-induced bodies (RLBs), which form upon the initiation of RNase L-mediated mRNA decay in response to viral infection and contain host/viral 3’-end RNA degradation intermediates (47–49). Because we observed these small punctate nsp1-dependent granules in RNase L KO cells (Fig. 7A), we propose that these RLB-like structures assemble as a consequence of nsp1-dependent degradation of cellular mRNA.

We propose two mechanisms by which nsp1 limits SG assembly. First, nsp1 reduces the PERK-mediated ISR activation (Fig. 3), which is consistent with observations made by Dolliver et al. (33). Second, our data indicate that the SGs are composed of cellular mRNA as opposed to viral ssRNA or dsRNA (Fig. 7A,D,E). This argues that these are not aggregated RNA condensates (VARCs), which form via G3BP1-mediated condensation of viral RNA (29). Notably, the small punctate nsp1-dependent granules in cells inrfected with WT SARS-CoV2 and the large TIA1/poly(A)+RNA-positivie SGs in cells infected with the nsp1 mutant SARS-CoV-2 lacked G3BP1 (Fig. 7A). We provide two explanations to account for these observations. First, RLBs, which are similar to the small punctate nsp1-dependent bodies, assemble in a G3BP1-independent manner and form during WT SARS-CoV-2 infection depite N-mediated inhibiton of G3BP1 (50). Thus, the nsp1-mediated RNA decay granules for independently of G3BP1. Second, in the cells infected with the nsp1 mutant viruses, we propose that the combined viral and host RNA loads in cells infected with nsp1 mutant SARS-CoV-2 drives the assembly of SGs despite N-mediated antagonism of G3BP1. These observations establish a function for nsp1-mediated decay of cellular mRNA in limiting antiviral SG assembly. Future studies will further investigate the role of the PERK-mediated ISR, G3BP1, GADD34, and cellular RNA decay in regulating SGs triggered by SARS-CoV-2 nsp1 mutants.

In summary, we provide important new evidence that details coronavirus-host competition over the translation machinery of the cell. Through the use of recombinant viruses, we demonstrate interactions between coronavirus nsp1 and the host ISR pathway. Moreover, we demonstrate differences between SARS-CoV-2 and MERS-CoV nsp1 translation inhibition and mRNA degradation that were not observed in overexpression and *in vitro* reconstitution studies. These studies provides a fundamental understanding of nsp1 that can be leveraged for drug discovery, live-attenuated vaccine development, and biotechnology applications.

## MATERIALS AND METHODS

All of the materials and methods are described in **SI Appendix, Materials & Methods**. Any materials or related protocols mentioned in this work can be obtained by contacting the corresponding authors.

## Acknowledgments

We thank Dr. Shinji Makino for sharing MERS-CoV mutant viruses and Dr. Mark Denison for sharing anti-MERS-CoV nsp1 rabbit antiserum. This work was supported by grants from the National Institutes of Health, R01AI140442 (SRW), R01AI169537 (SRW&NAC), R35GM151249 (JMB), R01A1AI161175 (LM-S), R01AI161363 (LM-S), R35GM138029 (ARF) P20GM113117 (ARF), Penn Center for Research on Emerging Viruses (SRW), a pilot grant from the Institute for RNA Innovation of the Perelman School of Medicine, the University of Pennsylvania, Herbert Wertheim University of Florida Scripps Institute for Biomedical Innovation and Technology (JMB). D.M.R. was supported in part by NIH T32 AI055400. J.J.P. was supported by a fellowship from the Madison and Lila Self graduate fellowship program at the University of Kansas.

## Supporting Information for

### This PDF file includes

Materials & Methods

Table S1

SI References

Figures S1 to S5 and legends

## MATERIALS AND METHODS

### Nasal cultures

Nasal epithelial cells were collected via cytologic brushing of patients’ nasal cavities after obtaining informed consent and then grown and differentiated on transwell inserts to establish ALI cultures. The full study protocol was approved by the University of Pennsylvania Institutional Review Board (protocol # 800614) and the Philadelphia VA Institutional Review Board (protocol #00781).

### Cell culture

VeroE6 cells and VeroCCL81 cells from American Type Culture Collection (ATCC) were cultured in Dulbecco’s modified Eagle’s medium (DMEM) containing 10% L-glutamine and 4.5g/L D-glucose (Gibco, ThermoFisher) supplemented with 10% heat inactivated (HI) fetal bovine serum (FBS) (Hyclone, Cytiva) and 1X penicillin/streptomycin (pen/strep) (Gibco, ThermoFisher). Human A549 cells engineered to stably express angiotensin-converting enzyme 2 (ACE2) (A549-ACE2) or dipeptidyl peptidase-4 (DPP4) (A549-DPP4) and the various CRISPR-Cas9 knock-out cell lines were cultured in RPMI 1640 (Gibco, ThermoFisher) supplemented with 10% HI FBS and 1X pen/strep.

### CRISPR/Cas9 engineered cell lines

*PKR* KO A549^ACE2^ and A549^DPP4^ cells were constructed using the same lenti-CRISPR system and guide sequences as previously described (1–3), but subjected to blasticidin (10ug/mL) selection. *PERK* KO A549^ACE2^ and A549^DPP4^ cells were constructed using the same lenti-CRISPR system and guide sequences as previously described (1–3), but subjected to blasticidin (10ug/mL) selection and with the following guide RNA: (5’-CACCGGGTACTCGCGTCGCTGAGG T – 3’). *RNASEL* KO A549^ACE2^ and A549^DPP4^ cells were generated previously (3, 4).

### Recombinant viruses

Recombinant SARS-CoV-2 (USA-WA1/2020 strain) viruses were derived from a bacterial artificial chromosome (BAC) vector containing the full-length SARS-CoV-2 USA-WA1/2020 genome (5). For the SARS-COV-2 nsp1 K164A/H165A mutant virus, nucleotides 755 (A), 756 (A), 758 (C), and 759 (A) were mutated in order to substitute both lysine 164 and histidine 165 with alanine residues. For the SARS-COV-2 nsp1 R124A/K125A mutant virus, nucleotides 635 (C), 636 (G), 638 (A), and 639 (A) were mutated in order to substitute both arginine 124 and lysine 125 with alanine residues. The SARS-COV-2 nsp1 R124A/K125A/K164A/H165A double mutant virus was made by combining both sets of mutations. All mutant viruses were rescued as previously described (5, 6). The SARS-CoV-2 mutant virus containing the D33K/E36K/E37K/E41L substitutions in nsp1 was previously described (7) and obtained from Dr. Luis Martinez-Sobrido. MERS-CoV (HCoV EMC/2012) nsp1 mutant viruses were previously described in (8). Virus stocks were sequenced and compared to the publicly available wild-type sequences on NCBI. All virus stocks were generated via low MOI infections in VeroE6 cells (for SARS-CoV-2 viruses) and VeroCCL81 cells (for MERS-CoV viruses). MERS-CoV viruses were obtained from Dr. Shinji Makino (8, 9).

### Infection

A549^ACE2^, A549^DPP4^, and primary nasal epithelial cells were infected with recombinant SARS-CoV-2 WT, SARS-CoV-2 nsp1 mutant, MERS-CoV WT, and MERS-CoV nsp1 mutant viruses at multiplicity of infection (MOI) of 1. At indicated times post infection, infectious virus in cellular supernatants or apical surface liquid collected via apical wash of nasal ALI cultures were quantified via plaque assay (10). Whole-cell lysates were also collected from infected cells for downstream Western blot analysis (11).

### RiboPuromycylation assays

RiboPuromycylation assays were conducted as previously described (12). Briefly, cells were infected with MOI=1 of each virus and, at 48hpi, treated with 10ug/mL puromycin, incubated at 37C for ten minutes, and whole cell lysates collected for Western blot analysis whereby an anti-puromycin antibody was used to measure puromycylation of proteins as a proxy for global translation by Western blot as below (13).

### Western blot

Cells were infected at MOI=1 for all infections and whole-cell lysates were collected at 48hpi. Whole-cell lysates were separated on 4-15% denaturing SDS-PAGE gels, transferred to PVDF membranes, blocked with 5% milk in 1X TBST for at least one hour, probed with primary antibodies at 4C overnight, and probed with HRP-conjugated secondary antibodies for 30-60 minutes before development and imaging (11).

### smFISH and immunofluorescence assays

smFISH was performed as previously described (14, 15). Oligos used to generate smFISH probes can be found in data file S1. Poly(A)+ RNA was detected using oligo(dT)_18_ Cy5 (Integrated DNA technologies). For immunofluorescence assays, cells were incubated with rabbit polyclonal anti-GADD34 antibody (1:1000; proteintech, 10449-1-AP), mouse monoclonal anti-G3BP1 antibody (1:1000; Abcam, ab119236), mouse monoclonal dsRNA antibody (K1) (1:1000; Cell Signaling Technology, 28764), rabbit recombinant monoclonal anti-TIA1 antibody (1:1000, Abcam, ab263945) for 4-8 hours. Cells were washed three times with phosphate-buffered saline (PBS) and incubated with goat anti-rabbit immunoglobulin G (IgG; Alexa Fluor 488) (Abcam, ab150077), goat anti-rabbit immunoglobulin G (IgG; Alexa Fluor 555) (Abcam, ab150078), or goat anti-mouse IgG (FITC) (Abcam; ab97022) at 1:1000 for 2 hours. After three washes, cells were either mounted for imaging or fixed with 4% PFA followed by the smFISH protocol.

Microscopy was performed as previously described (14). Coverslips were mounted on slides with VECTASHIELD Antifade Mounting Medium with 4′,6-diamidino-2-phenylindole (DAPI; Vector Laboratories; H-1200). A Nikon Eclipse Ti2 equipped with a Yokogawa CSU-W1 spinning disk confocal, 100×, 1.45 NA oil objective was used for imaging with Nikon NIS-Elements software. Images were processed using ImageJ with the FIJI plugin. *Z* planes were stacked (Max intensity display), and minimum and maximum display values were set in ImageJ for each channel to properly view fluorescence. Fluorescence intensity and smFISH foci were quantified in ImageJ using “measure” and “analyze particles” tools, respectively.

### Statistical analysis

Image processing was conducted in Fiji (ImageJ2) and data processing was conducted in Microsoft Excel. Data was graphed and statistics were performed using GraphPad Prism. Data was graphed displaying either individual values or the mean +/− standard deviation. For viral growth curves (Fig 2), statistical significance was determined by comparing mutant viruses to WT using a two-way ANOVA for experiments with multiple time points and/or viruses. For IFA and smFISH, statistical significance was determined by comparing WT to individual mutants using multiple t-tests with multiple comparison correction (Benjamini and Hochberg) and validated using two-way ANOVA. Significance shown represents P values, where * = P < 0.05; ** = P < 0.01; *** = P < 0.001; and **** = P < 0.0001. Nonsignificant comparisons are not displayed in all figures.

**Table S1.**
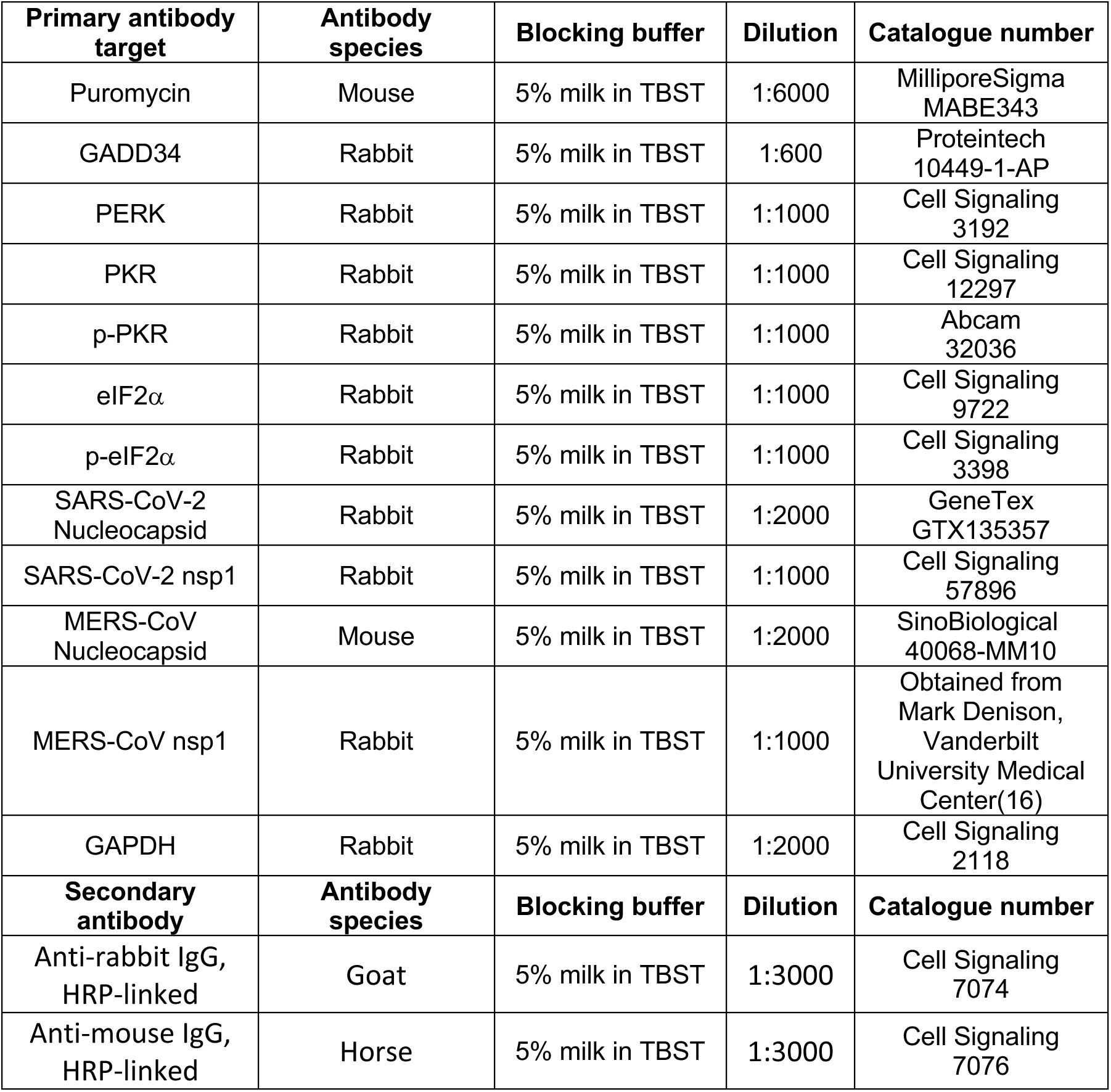
Antibodies.

---

**Figure S1.**
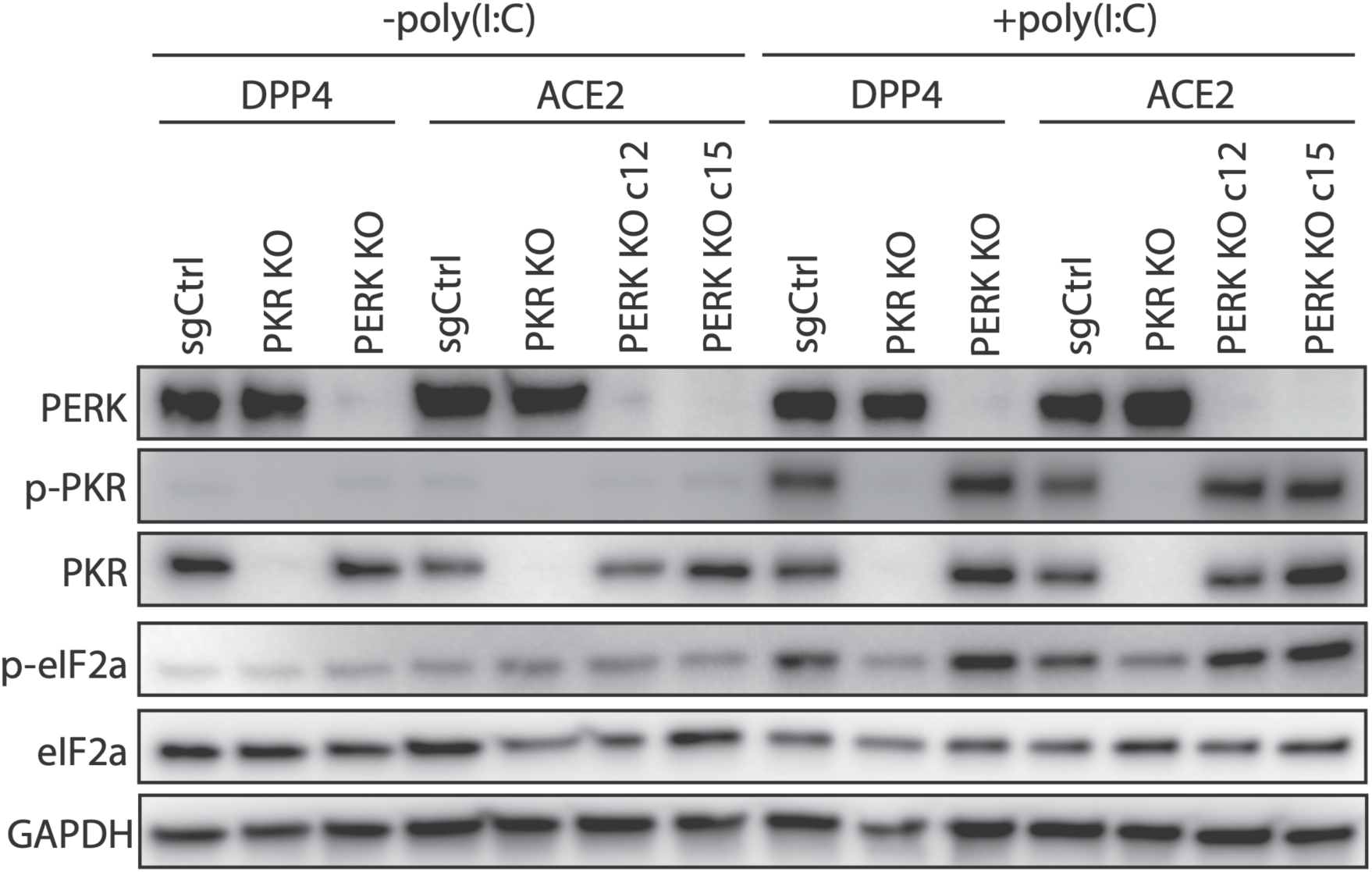
Protein expression in poly(I:C) transfected cells. Indicated cells were transfected with 200ng of high-molecular weight (HMW) poly(I:C) (+poly(I:C)) or scrambled siRNA control (-poly(I:C)) for 4 hours. Whole-cell lysates were collected at 4 hours post-transfection, analyzed by western blot, and probed with antibodies as indicated. Cell receptors, KO, and PERK KO clone c12 or c15) are indicated. Data shown are from one representative of two independent experiments.

**Figure S2.**
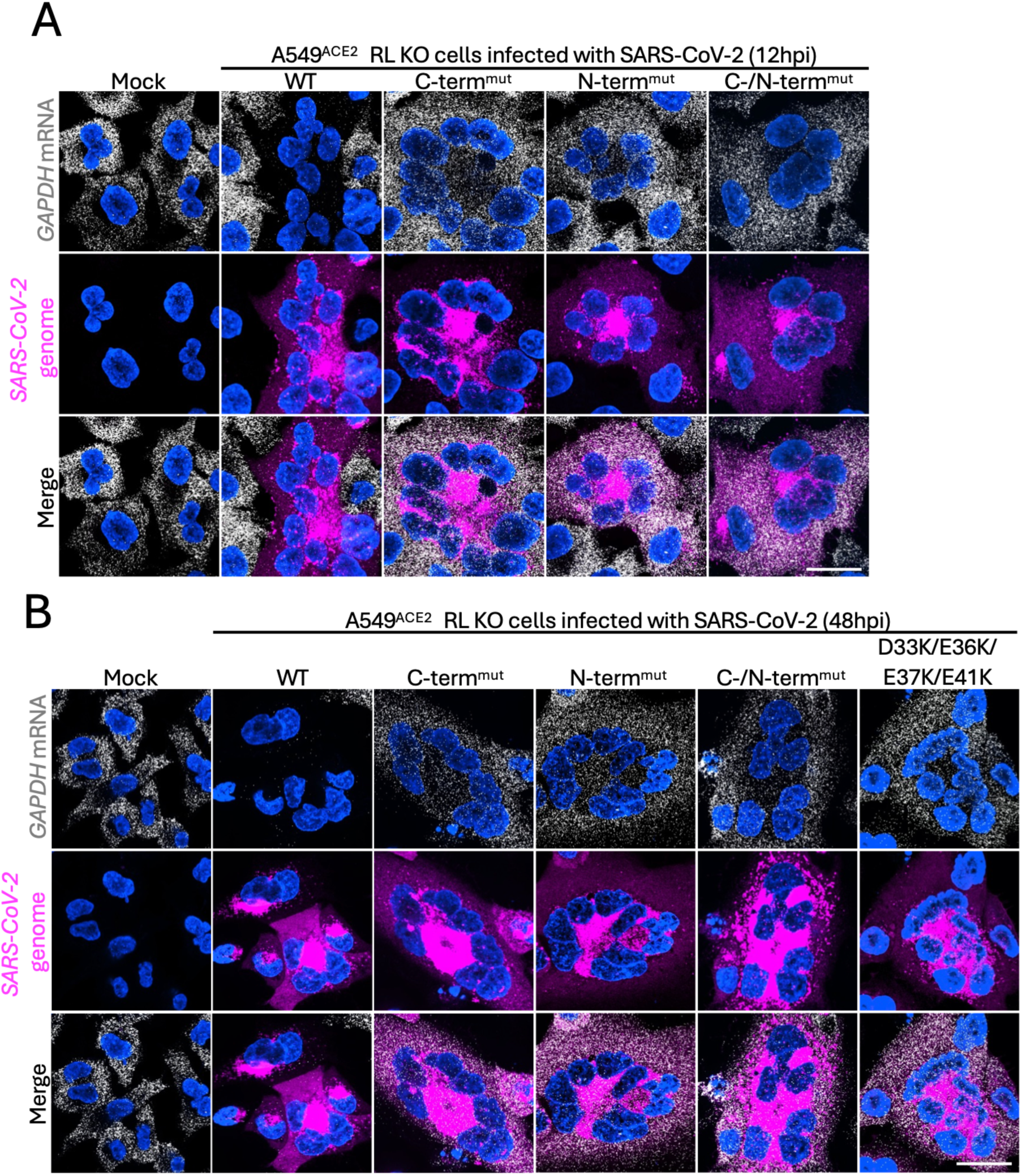
SARS-CoV-2 and MERS-CoV nsp1 is required for viral-mediated degradation of cellular mRNA. smFISH for SARS-CoV-2 genome RNA (ORF1a) and human *GAPDH* mRNA in A549^ACE2^ RL KO cells infected with WT or nsp1 mutant SARS-CoV-2 viruses at 12hpi (A) and 48hpi (B). Images are representative of 3 biological replicates. Scale bar represents 20μm.

**Figure S3.**
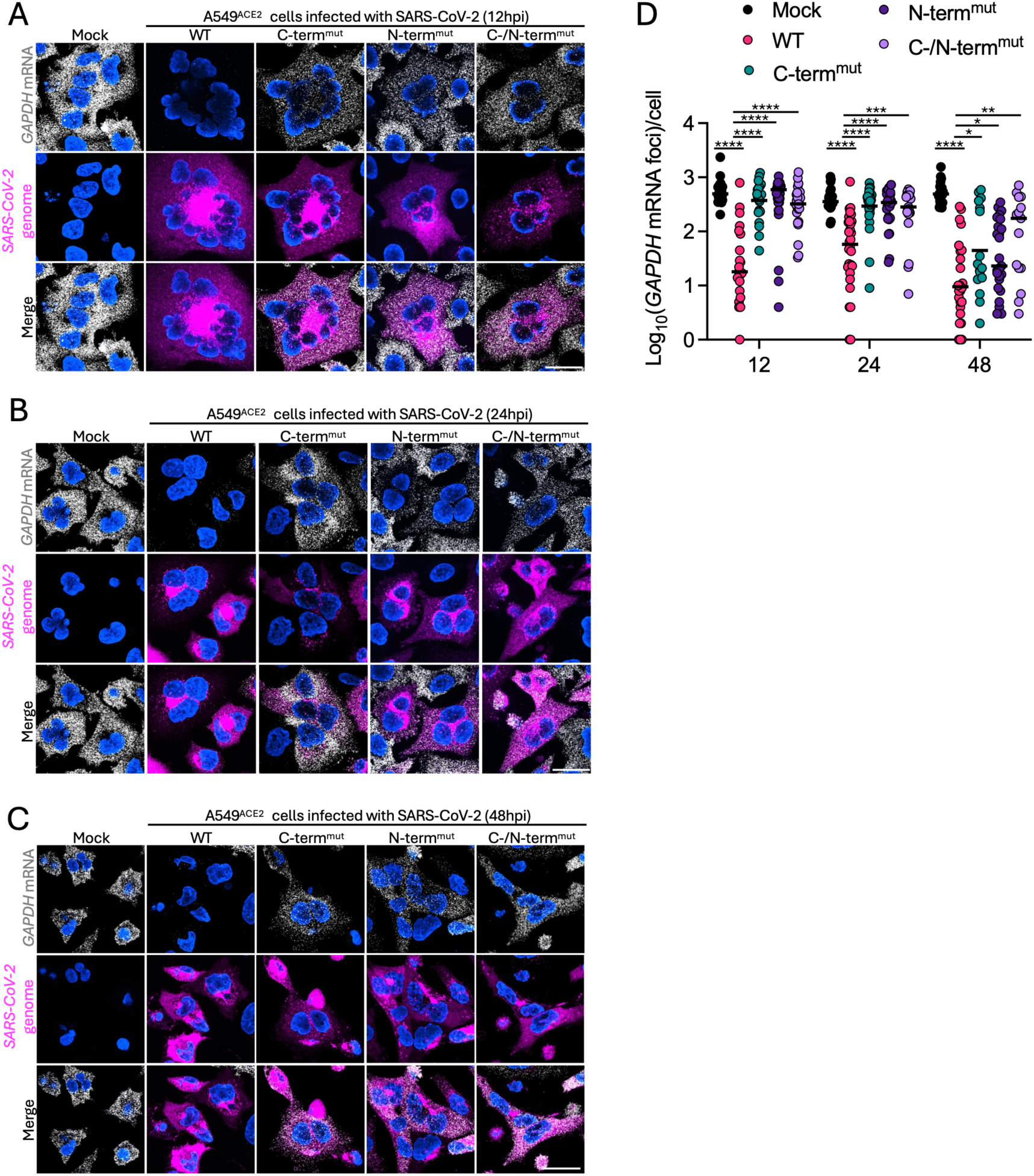
SARS-CoV-2 and MERS-CoV nsp1 is required for viral-mediated degradation of cellular mRNA. smFISH for SARS-CoV-2 genome RNA (ORF1a) and human *GAPDH* mRNA in A549^ACE2^ cells infected with WT or nsp1 mutant SARS-CoV-2 viruses at 12 hpi (A), 24hpi (B), and 48hpi (C). Images are representative of 3 biological replicates. (D) Quantification of *GAPDH* mRNA in individual cells (dots) at indicated time post-infection as represented in (A-C). Minimum of 15 cells per virus were quantified. Quantification is representative of 3 biological replicates. Scale bar represents 20μm.

**Figure S4.**
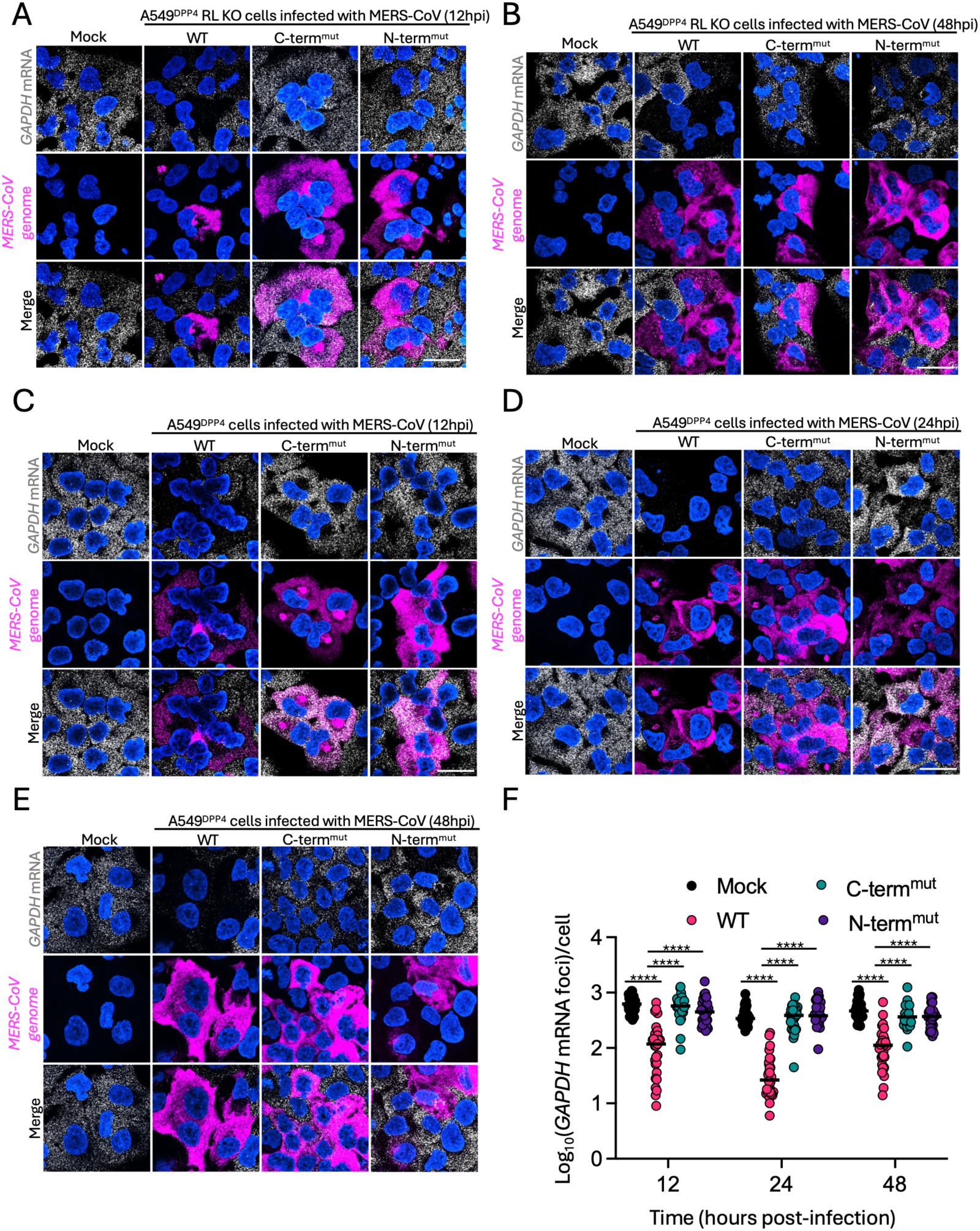
SARS-CoV-2 and MERS-CoV nsp1 is required for viral-mediated degradation of cellular mRNA. smFISH for MERS-CoV genome RNA (ORF1a) and human *GAPDH* mRNA in A549^DPP4^ RL KO cells infected with WT or nsp1 mutant MERS-CoV viruses at 12hpi (A) and 48hpi (B). Images are representative of 3 biological replicates. smFISH for MERS-CoV genome RNA (ORF1a) and human *GAPDH* mRNA in A549^DPP4^ cells infected with WT or nsp1 mutant MERS-CoV viruses at 12hpi (C), 24hpi (D), and 48hpi (E). Images are represenetative of 3 biological replicates. (F) Quantification of *GAPDH* mRNA in individual cells (dots) at indicated time post-infection as represented in (C-E). Minimum of 23 cells per virus was quantified. Quantification is representative of 3 biological replicates. Scale bar represents 20μm.

**Figure S5.**
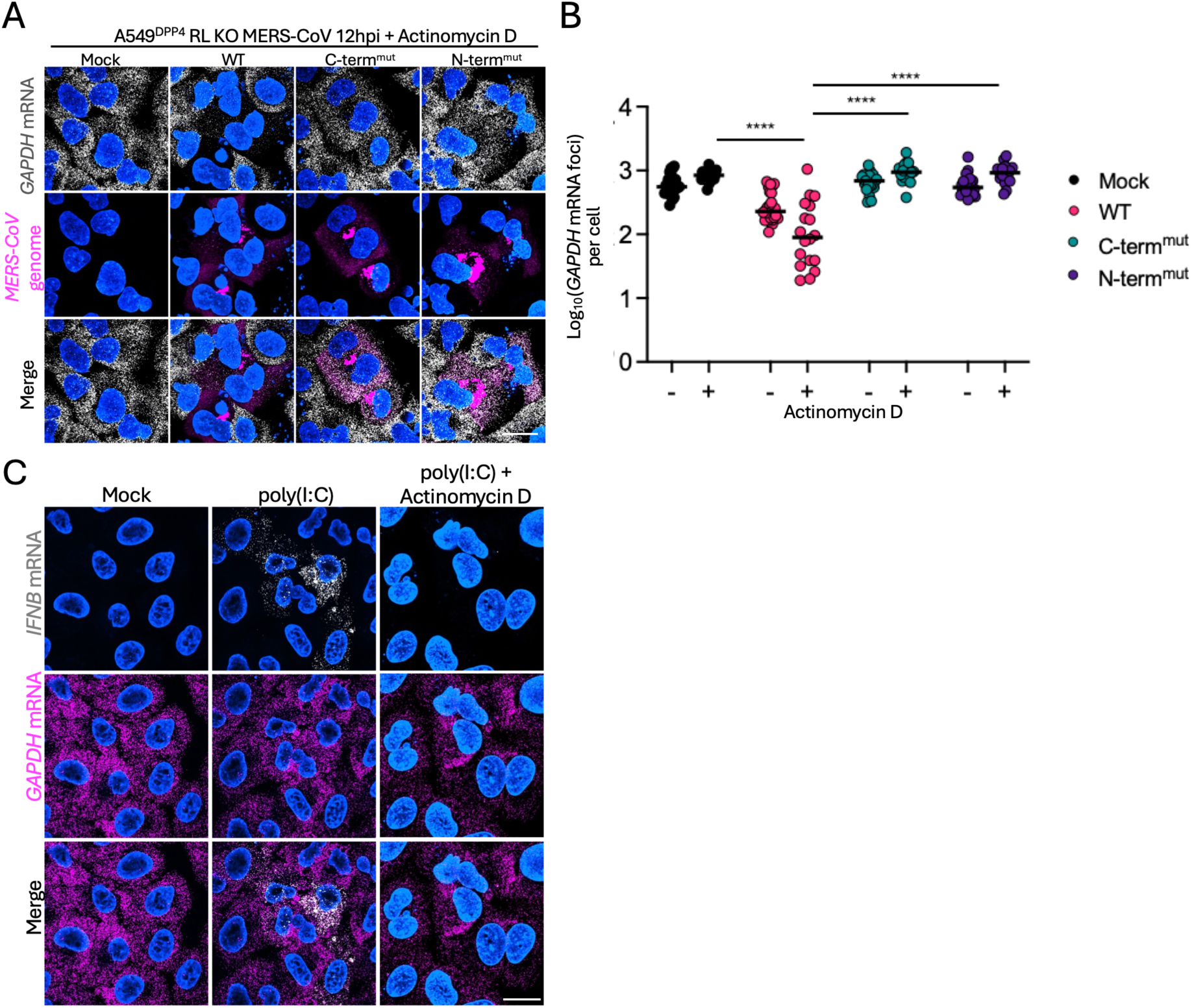
SARS-CoV-2 and MERS-CoV nsp1 is required for viral-mediated degradation of cellular mRNA. (A) smFISH for MERS-CoV genome RNA (ORF1a) and human *GAPDH* mRNA in A549^DPP4^ RL KO cells infected with WT or nsp1 mutant MERS-CoV viruses at 12hpi with 1mg/mL of Actinomycin D. Images are representative of 3 or more fields of view. (B) Quantification of *GAPDH* mRNA in individual cells (dots) for indicated viruses with or without 1mg/mL Actinomycin D treatment as represented in (A). Minimum of 16 cells per virus were quantified from 3 or more fields of view. (C) smFISH for human *GAPDH* mRNA and interferon beta (*IFNB*) mRNA in A549 RL KO cells transfected with 500ng of HMW poly(I:C) and co-treated with or without actinomycin D for 16 hours. Images aare representative of 3 or more fields of view. Scale bar represents 20μm.

## REFERENCES

1. K. Schubert et al., SARS-CoV-2 Nsp1 binds the ribosomal mRNA channel to inhibit translation. Nat Struct Mol Biol 27, 959–966 (2020).

2. M. Thoms et al., Structural basis for translational shutdown and immune evasion by the Nsp1 protein of SARS-CoV-2. Science 369, 1249–1255 (2020).

3. K. Schubert et al., Universal features of Nsp1-mediated translational shutdown by coronaviruses. Mol Cell 83, 3546–3557 e3548 (2023).

4. W. Kamitani, C. Huang, K. Narayanan, K. G. Lokugamage, S. Makino, A two-pronged strategy to suppress host protein synthesis by SARS coronavirus Nsp1 protein. Nat Struct Mol Biol 16, 1134–1140 (2009).

5. C. P. Lapointe et al., Dynamic competition between SARS-CoV-2 NSP1 and mRNA on the human ribosome inhibits translation initiation. Proc Natl Acad Sci U S A 118 (2021).

6. H. Kang, M. Feng, M. E. Schroeder, D. P. Giedroc, J. L. Leibowitz, Stem-loop 1 in the 5’ UTR of the SARS coronavirus can substitute for its counterpart in mouse hepatitis virus. Adv Exp Med Biol 581, 105–108 (2006).

7. L. Li et al., Structural lability in stem-loop 1 drives a 5’ UTR-3’ UTR interaction in coronavirus replication. J Mol Biol 377, 790–803 (2008).

8. P. Liu, D. Yang, K. Carter, F. Masud, J. L. Leibowitz, Functional analysis of the stem loop S3 and S4 structures in the coronavirus 3’UTR. Virology 443, 40–47 (2013).

9. S. Steiner et al., SARS-CoV-2 biology and host interactions. Nat Rev Microbiol 22, 206–225 (2024).

10. K. G. Lokugamage, K. Narayanan, C. Huang, S. Makino, Severe acute respiratory syndrome coronavirus protein nsp1 is a novel eukaryotic translation inhibitor that represses multiple steps of translation initiation. J Virol 86, 13598–13608 (2012).

11. T. Tanaka, W. Kamitani, M. L. DeDiego, L. Enjuanes, Y. Matsuura, Severe acute respiratory syndrome coronavirus nsp1 facilitates efficient propagation in cells through a specific translational shutoff of host mRNA. J Virol 86, 11128–11137 (2012).

12. I. S. Abaeva, Y. Arhab, A. Miscicka, C. U. T. Hellen, T. V. Pestova, In vitro reconstitution of SARS-CoV-2 Nsp1-induced mRNA cleavage reveals the key roles of the N-terminal domain of Nsp1 and the RRM domain of eIF3g. Genes Dev 37, 844–860 (2023).

13. Y. Tardivat et al., SARS-CoV-2 NSP1 induces mRNA cleavages on the ribosome. Nucleic Acids Res 51, 8677–8690 (2023).

14. J. M. Burke, L. A. St Clair, R. Perera, R. Parker, SARS-CoV-2 infection triggers widespread host mRNA decay leading to an mRNA export block. RNA 27, 1318–1329 (2021).

15. K. Nakagawa et al., The Endonucleolytic RNA Cleavage Function of nsp1 of Middle East Respiratory Syndrome Coronavirus Promotes the Production of Infectious Virus Particles in Specific Human Cell Lines. J Virol 92 (2018).

16. K. G. Lokugamage et al., Middle East Respiratory Syndrome Coronavirus nsp1 Inhibits Host Gene Expression by Selectively Targeting mRNAs Transcribed in the Nucleus while Sparing mRNAs of Cytoplasmic Origin. J Virol 89, 10970–10981 (2015).

17. T. Fisher et al., Parsing the role of NSP1 in SARS-CoV-2 infection. bioRxiv 10.1101/2022.03.14.484208 (2022).

18. K. Narayanan et al., Severe acute respiratory syndrome coronavirus nsp1 suppresses host gene expression, including that of type I interferon, in infected cells. J Virol 82, 4471–4479 (2008).

19. S. I. Shehata, R. Parker, SARS-CoV-2 Nsp1 mediated mRNA degradation requires mRNA interaction with the ribosome. RNA Biol 20, 444–456 (2023).

20. Y. Li et al., SARS-CoV-2 induces double-stranded RNA-mediated innate immune responses in respiratory epithelial-derived cells and cardiomyocytes. Proc Natl Acad Sci U S A 118 (2021).

21. D. M. Renner, N. A. Parenti, N. Bracci, S. R. Weiss, Betacoronaviruses Differentially Activate the Integrated Stress Response to Optimize Viral Replication in Lung-Derived Cell Lines. Viruses 17 (2025).

22. C. E. Comar et al., MERS-CoV endoribonuclease and accessory proteins jointly evade host innate immunity during infection of lung and nasal epithelial cells. Proc Natl Acad Sci U S A 119, e2123208119 (2022).

23. C. J. Otter et al., SARS-CoV-2 nsp15 endoribonuclease antagonizes dsRNA-induced antiviral signaling. Proc Natl Acad Sci U S A 121, e2320194121 (2024).

24. D. M. Renner, N. A. Parenti, S. R. Weiss, Betacoronaviruses Differentially Activate the Integrated Stress Response to Optimize Viral Replication in Lung Derived Cell Lines. bioRxiv 10.1101/2024.09.25.614975, 2024.2009.2025.614975 (2024).

25. E. Kojima et al., The function of GADD34 is a recovery from a shutoff of protein synthesis induced by ER stress: elucidation by GADD34-deficient mice. FASEB J 17, 1573–1575 (2003).

26. Y. Y. Lee, R. C. Cevallos, E. Jan, An upstream open reading frame regulates translation of GADD34 during cellular stresses that induce eIF2alpha phosphorylation. J Biol Chem 284, 6661–6673 (2009).

27. S. Jain et al., ATPase-Modulated Stress Granules Contain a Diverse Proteome and Substructure. Cell 164, 487–498 (2016).

28. D. S. W. Protter, R. Parker, Principles and Properties of Stress Granules. Trends Cell Biol 26, 668–679 (2016).

29. J. M. Burke, O. C. Ratnayake, J. M. Watkins, R. Perera, R. Parker, G3BP1-dependent condensation of translationally inactive viral RNAs antagonizes infection. Sci Adv 10, eadk8152 (2024).

30. Z. Yang et al., Interaction between host G3BP and viral nucleocapsid protein regulates SARS-CoV-2 replication and pathogenicity. Cell Rep 43, 113965 (2024).

31. H. Liu et al., SARS-CoV-2 N Protein Antagonizes Stress Granule Assembly and IFN Production by Interacting with G3BPs to Facilitate Viral Replication. J Virol 96, e0041222 (2022).

32. M. Biswal, J. Lu, J. Song, SARS-CoV-2 Nucleocapsid Protein Targets a Conserved Surface Groove of the NTF2-like Domain of G3BP1. J Mol Biol 434, 167516 (2022).

33. S. M. Dolliver et al., Nsp1 proteins of human coronaviruses HCoV-OC43 and SARS-CoV2 inhibit stress granule formation. PLoS Pathog 18, e1011041 (2022).

34. C. J. Otter et al., Infection of primary nasal epithelial cells differentiates among lethal and seasonal human coronaviruses. Proc Natl Acad Sci U S A 120, e2218083120 (2023).

35. M. Mei et al., Inhibition of mRNA nuclear export promotes SARS-CoV-2 pathogenesis. Proc Natl Acad Sci U S A 121, e2314166121 (2024).

36. J. M. Burke, S. L. Moon, T. Matheny, R. Parker, RNase L Reprograms Translation by Widespread mRNA Turnover Escaped by Antiviral mRNAs. Mol Cell 75, 1203–1217 e1205 (2019).

37. A. R. Jauregui, D. Savalia, V. K. Lowry, C. M. Farrell, M. G. Wathelet, Identification of residues of SARS-CoV nsp1 that differentially affect inhibition of gene expression and antiviral signaling. PLoS One 8, e62416 (2013).

38. M. G. Wathelet, M. Orr, M. B. Frieman, R. S. Baric, Severe acute respiratory syndrome coronavirus evades antiviral signaling: role of nsp1 and rational design of an attenuated strain. J Virol 81, 11620–11633 (2007).

39. I. Frolov et al., All Domains of SARS-CoV-2 nsp1 Determine Translational Shutoff and Cytotoxicity of the Protein. J Virol 97, e0186522 (2023).

40. S. Yuan et al., Nonstructural Protein 1 of SARS-CoV-2 Is a Potent Pathogenicity Factor Redirecting Host Protein Synthesis Machinery toward Viral RNA. Mol Cell 80, 1055–1066 e1056 (2020).

41. A. K. Banerjee et al., SARS-CoV-2 Disrupts Splicing, Translation, and Protein Trafficking to Suppress Host Defenses. Cell 183, 1325–1339 e1321 (2020).

42. A. S. Mendez et al., The N-terminal domain of SARS-CoV-2 nsp1 plays key roles in suppression of cellular gene expression and preservation of viral gene expression. Cell Rep 37, 109841 (2021).

43. C. E. Comar et al., Antagonism of dsRNA-Induced Innate Immune Pathways by NS4a and NS4b Accessory Proteins during MERS Coronavirus Infection. mBio 10 (2019).

44. J. J. Berlanga et al., The differential effect of SARS-CoV-2 NSP1 on mRNA translation and stability reveals new insights linking ribosome recruitment, codon usage, and virus evolution. Nucleic Acids Res 53 (2025).

45. V. Magg et al., Turnover of PPP1R15A mRNA encoding GADD34 controls responsiveness and adaptation to cellular stress. Cell Rep 43, 114069 (2024).

46. S. Rao et al., Genes with 5’ terminal oligopyrimidine tracts preferentially escape global suppression of translation by the SARS-CoV-2 Nsp1 protein. RNA 27, 1025–1045 (2021).

47. J. M. Burke, E. T. Lester, D. Tauber, R. Parker, RNase L promotes the formation of unique ribonucleoprotein granules distinct from stress granules. J Biol Chem 295, 1426–1438 (2020).

48. J. M. Watkins, J. M. Burke, A closer look at mammalian antiviral condensates. Biochem Soc Trans 52, 1393–1404 (2024).

49. J. M. Watkins, J. M. Burke, RNase L-induced bodies sequester subgenomic flavivirus RNAs to promote viral RNA decay. Cell Rep 43, 114694 (2024).

50. J. M. Burke, Regulation of ribonucleoprotein condensates by RNase L during viral infection. Wiley Interdiscip Rev RNA 14, e1770 (2023).

51. L. Zhao et al., Antagonism of the interferon-induced OAS-RNase L pathway by murine coronavirus ns2 protein is required for virus replication and liver pathology. Cell Host Microbe 11, 607–616 (2012).

52. Y. Li et al., Activation of RNase L is dependent on OAS3 expression during infection with diverse human viruses. Proc Natl Acad Sci U S A 113, 2241–2246 (2016).

53. C. Ye et al., Rescue of SARS-CoV-2 from a Single Bacterial Artificial Chromosome. mBio 11 (2020).

54. C. Ye, L. Martinez-Sobrido, Use of a Bacterial Artificial Chromosome to Generate Recombinant SARS-CoV-2 Expressing Robust Levels of Reporter Genes. Microbiol Spectr 10, e0273222 (2022).

55. M. Piecyk, J. Fauvre, C. Duret, C. Chaveroux, C. Ferraro-Peyret, SUrface SEnsing of Translation (SUnSET), a Method Based on Western Blot Assessing Protein Synthesis Rates in vitro. Bio Protoc 14, e4933 (2024).

56. R. Cusic, J. M. Burke, Condensation of RNase L promotes its rapid activation in response to viral infection in mammalian cells. Sci Signal 17, eadi9844 (2024).

57. R. Cusic, J. M. Watkins, J. M. Burke, Single-cell analysis of RNase L-mediated mRNA decay. Methods Enzymol 692, 157–175 (2023).

## REFERENCES

1. L. Zhao et al., Antagonism of the interferon-induced OAS-RNase L pathway by murine coronavirus ns2 protein is required for virus replication and liver pathology. Cell Host Microbe 11, 607–616 (2012).

2. Y. Li et al., Activation of RNase L is dependent on OAS3 expression during infection with diverse human viruses. Proc Natl Acad Sci U S A 113, 2241–2246 (2016).

3. C. E. Comar et al., Antagonism of dsRNA-Induced Innate Immune Pathways by NS4a and NS4b Accessory Proteins during MERS Coronavirus Infection. mBio 10 (2019).

4. J. M. Burke, L. A. St Clair, R. Perera, R. Parker, SARS-CoV-2 infection triggers widespread host mRNA decay leading to an mRNA export block. RNA 27, 1318–1329 (2021).

5. C. Ye et al., Rescue of SARS-CoV-2 from a Single Bacterial Artificial Chromosome. mBio 11 (2020).

6. C. Ye, L. Martinez-Sobrido, Use of a Bacterial Artificial Chromosome to Generate Recombinant SARS-CoV-2 Expressing Robust Levels of Reporter Genes. Microbiol Spectr 10, e0273222 (2022).

7. M. Mei et al., Inhibition of mRNA nuclear export promotes SARS-CoV-2 pathogenesis. Proc Natl Acad Sci U S A 121, e2314166121 (2024).

8. K. G. Lokugamage et al., Middle East Respiratory Syndrome Coronavirus nsp1 Inhibits Host Gene Expression by Selectively Targeting mRNAs Transcribed in the Nucleus while Sparing mRNAs of Cytoplasmic Origin. J Virol 89, 10970–10981 (2015).

9. K. Nakagawa et al., The Endonucleolytic RNA Cleavage Function of nsp1 of Middle East Respiratory Syndrome Coronavirus Promotes the Production of Infectious Virus Particles in Specific Human Cell Lines. J Virol 92 (2018).

10. C. J. Otter et al., Infection of primary nasal epithelial cells differentiates among lethal and seasonal human coronaviruses. Proc Natl Acad Sci U S A 120, e2218083120 (2023).

11. Y. Li et al., SARS-CoV-2 induces double-stranded RNA-mediated innate immune responses in respiratory epithelial-derived cells and cardiomyocytes. Proc Natl Acad Sci U S A 118 (2021).

12. M. Piecyk, J. Fauvre, C. Duret, C. Chaveroux, C. Ferraro-Peyret, SUrface SEnsing of Translation (SUnSET), a Method Based on Western Blot Assessing Protein Synthesis Rates in vitro. Bio Protoc 14, e4933 (2024).

13. D. M. Renner, N. A. Parenti, N. Bracci, S. R. Weiss, Betacoronaviruses Differentially Activate the Integrated Stress Response to Optimize Viral Replication in Lung-Derived Cell Lines. Viruses 17 (2025).

14. R. Cusic, J. M. Burke, Condensation of RNase L promotes its rapid activation in response to viral infection in mammalian cells. Sci Signal 17, eadi9844 (2024).

15. R. Cusic, J. M. Watkins, J. M. Burke, Single-cell analysis of RNase L-mediated mRNA decay. Methods Enzymol 692, 157–175 (2023).

16. E. Prentice, J. McAuliffe, X. Lu, K. Subbarao, M. R. Denison, Identification and characterization of severe acute respiratory syndrome coronavirus replicase proteins. J Virol 78, 9977–9986 (2004).

